# Structural insights into transcriptional regulation by the helicase RECQL5

**DOI:** 10.1101/2025.01.29.634372

**Authors:** Alfredo Jose Florez Ariza, Nicholas Z. Lue, Patricia Grob, Benjamin Kaeser, Jie Fang, Susanne A. Kassube, Eva Nogales

## Abstract

Transcription and its regulation pose a major challenge for genome stability. The helicase RECQL5 has been proposed as an important factor to help safeguard the genome, and is the only member of the human RecQ helicase family that directly binds to RNA Polymerase II (Pol II) and affects its progression. RECQL5 mitigates transcription stress and genome instability in cells, yet the molecular mechanism underlying this phenomenon is unclear. Here, we employ cryo-electron microscopy (cryo-EM) to determine the structures of stalled Pol II elongation complexes (ECs) bound to RECQL5. Our structures reveal the molecular interactions stabilizing RECQL5 binding to the Pol II EC and highlight its role as a transcriptional roadblock. Additionally, we find that RECQL5 can modulate the Pol II translocation state. In its nucleotide-free state, RECQL5 mechanically twists the downstream DNA in the EC, and upon nucleotide binding, it undergoes a conformational change that allosterically induces Pol II towards a post-translocation state. We propose this mechanism may help restart Pol II elongation and therefore contribute to reduction of transcription stress.

## Introduction

Despite its fundamental importance for life, transcription has genome destabilizing effects that pose challenges for cells^1^. Collisions between transcribing RNA polymerases and the replication machinery, for example, can stall replication forks and are well-known sources of DNA damage^1–3^. Cells therefore require mechanisms to resolve such conflicts and mitigate the deleterious side effects of transcription. RecQ helicases are key players in these efforts to safeguard genome stability. In humans, the RecQ family comprises five 3’ to 5’ DNA helicases: RECQL1, BLM (Bloom syndrome helicase), WRN (Werner syndrome helicase), RECQL4, and RECQL5^3,4^. Testifying to the importance of this family, mutations in several RecQ genes—*BLM*, *WRN*, and *RECQL4*—are associated with human genetic disorders marked at a cellular level by erosion of genome integrity^3–7^. Although RECQL5 has not been conclusively linked to human diseases, loss of RECQL5 is known to promote cancer in mice, likely as a result of increased genome instability^8^. Consistently, there is some evidence linking RECQL5 loss-of-function to breast cancer susceptibility in humans^9^. On the other hand, RECQL5 has also been shown to be important for triple-negative breast cancer cell growth, as it counters excessive DNA damage arising from replication stress in these cells^10^. These findings underscore the importance of RECQL5’s multifaceted roles within the cell.

RECQL5 has been proposed to protect cells against genome instability through several mechanisms. It has long been known that RECQL5 can remove filaments of RAD51 from single-stranded DNA through its helicase activity, thereby suppressing inappropriate homologous recombination^8,11,12^. This disassembly of RAD51 filaments is also a crucial step in the resolution of stalled replication forks and is important for proper replication restart after transcription-replication conflicts^13–15^. Among the RecQ helicases, RECQL5 is also unique in that it directly interacts with Pol II^16–20^. Through this interaction, RECQL5 inhibits transcription both in vitro and in cells^17,21–23^. In cells, loss of RECQL5 has been shown to promote faster transcription but with both greater transcription stress (Pol II pausing, stalling, and arrest) and increased rates of chromosomal rearrangements^23^. These results indicate that RECQL5 enhances transcriptional robustness, but the mechanistic basis of this function has not yet been elucidated.

In our previous work, we showed that RECQL5’s internal Pol II-interacting (IRI) module, which consists of the αN helix and KIX domain (**Fig. 1a**), is critical for its interaction with Pol II^22^. This work revealed that RECQL5’s KIX domain competes with the transcription elongation factor TFIIS for interaction with Pol II’s RPB1 subunit, thereby inhibiting Pol II progression through pause sites. Our low-resolution structure (∼13 Å) also showed that RECQL5’s helicase domain binds to downstream DNA but was insufficient to define the molecular interactions underlying RECQL5’s ability to serve as a roadblock for Pol II advancement. In the decade since this initial study, no other structures have been reported on RECQL5 in the context of transcriptional regulation, emphasizing the need for further detailed structural study.

**Fig. 1.**
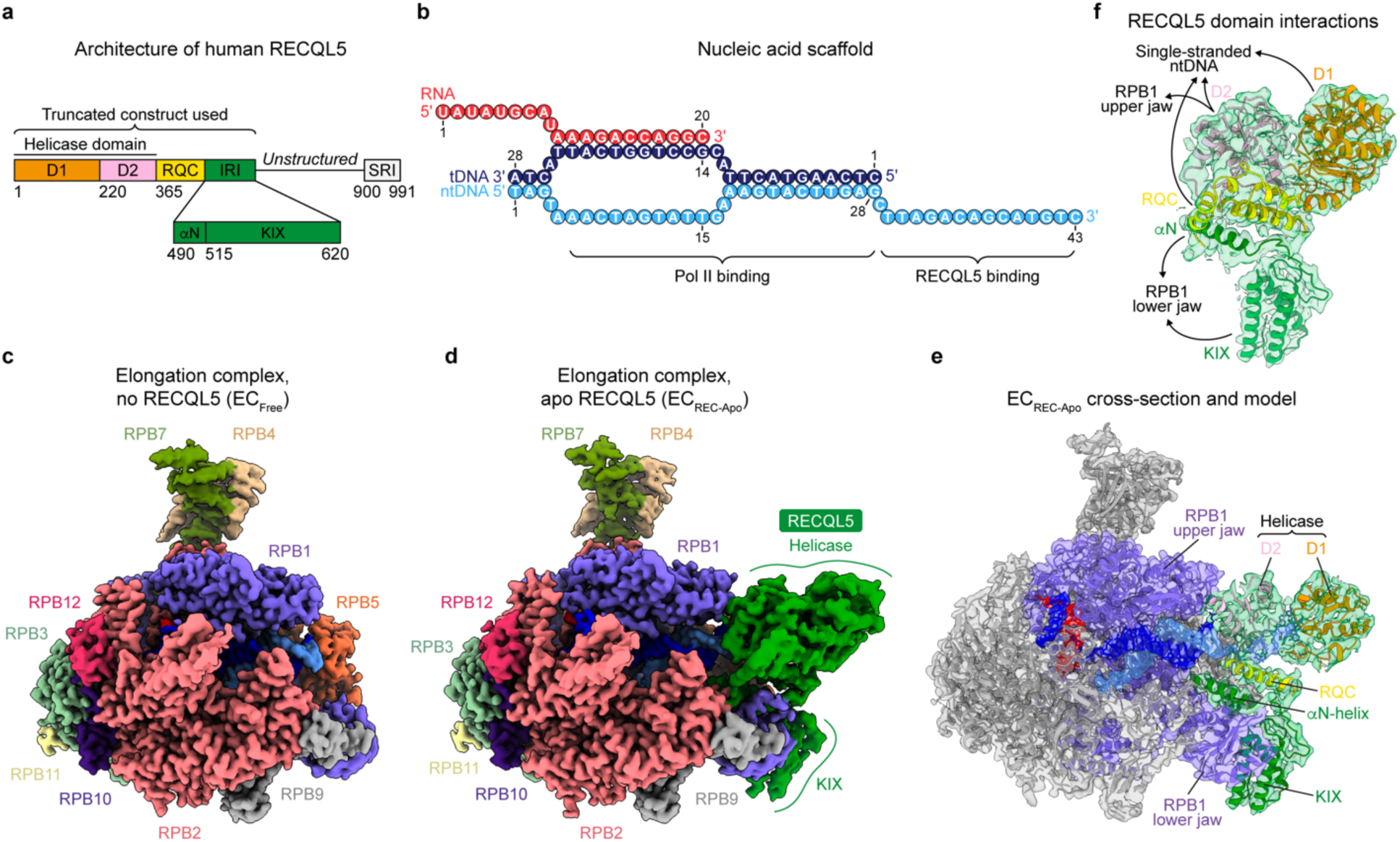
| Structure of the Pol II EC bound to RECQL5. **a**. Domain architecture of RECQL5 indicating the truncated construct used in this study (residues 1–620). The helicase domain (comprising the D1 and D2 subdomains), RQC (RecQ C-terminal) domain, IRI (Internal Pol II-interacting) module (comprising the αN helix and KIX domain), and SRI (Set2-Rpb1-interacting) domain are indicated. **b**. Schematic depicting the DNA/RNA scaffold used in this study. Numbers indicate nucleotide position along each strand. **c**. Cryo-EM map of free Pol II EC (EC_Free_). Pol II subunits are colored as indicated by the labels, and nucleic acids are colored as in **b**. **d**. Cryo-EM map of Pol II EC bound to RECQL5_1–620_ D157A (EC_REC-Apo_). Colors are the same as in **c,** with RECQL5 colored in green. **e**. Cross section of the EC_REC-Apo_ cryo-EM map (shown in transparency) with the fitted atomic model (shown in ribbon representation). Pol II is shown in gray apart from the RPB1 subunit shown in purple. The RECQL5 model is colored by domain as in **a** and the nucleic acids are colored as in **b**. **f**. View of the cryo-EM density for RECQL5 from EC_REC-Apo_ (shown in transparency) with the fitted atomic model (shown in ribbon representation), indicating the contacts each domain makes with the EC. Colors are the same as in **e**.

Understanding the molecular mechanism of RECQL5 transcriptional regulation requires a comprehension of the structural basis of Pol II transcription. During transcription, incorporation of a nucleotide into the growing RNA chain by Pol II generates an active site configuration referred to as a pre-translocation state^24^. The polymerase then translocates forward, generating a post-translocation state in which a new free binding site is available for the next incoming nucleotide. Multiple studies, employing both X-ray crystallography and cryo-EM, have elucidated the structures of Pol II in different translocation states^25–28^. Moreover, a crystallographic structure of Pol II bound to the transcriptional inhibitor α-amanitin showed that Pol II can also adopt a translocation intermediate conformation in addition to these two discrete structural states^29^.

Here, we use cryo-EM to determine the structures of Pol II ECs engaged with RECQL5 in order to gain molecular insight into RECQL5 transcription regulation. The improvement in resolution over our previous RECQL5–Pol II structure enables us to visualize the specific molecular contacts RECQL5 establishes with the Pol II EC. We find that RECQL5 has the ability to perturb Pol II’s translocation state, suggesting a mechanical role in helping restart stalled Pol II.

## Results

### Structure of the Pol II EC bound to RECQL5

To study the molecular basis for RECQL5’s inhibitory effect on transcription, we conducted single-particle cryo-EM analysis of an in vitro reconstituted stalled human Pol II EC bound to RECQL5. The EC was assembled using a nucleic acid scaffold comprising both template and non-template strand DNAs (tDNA and ntDNA, respectively) as well as a hybridized RNA, mimicking a transcription bubble^22,25^ (**Fig. 1b**). Additionally, the scaffold contained an extended single-stranded DNA (ssDNA) region (3’ end of the ntDNA) around the Pol II DNA entry site. This scaffold provided a platform for RECQL5 to bind in a head-to-head orientation with respect to Pol II^22^. For our structural studies, we employed a truncated RECQL5 construct (RECQL5_1–620_) encompassing the helicase domain (D1 and D2 subdomains), RecQ C-terminal (RQC) domain, and the IRI module (comprising the αN helix and KIX domain) (**Fig. 1a**). RECQL5’s unstructured region and C-terminal Set2-Rpb1-interacting (SRI) domain were not included in this construct. Purified and mildly crosslinked EC bound to RECQL5 (catalytically inactive D157A mutant and without addition of ATP, EC_REC-Apo_) was deposited on graphene oxide grids and subjected to cryo-EM imaging.

Our cryo-EM processing showed the coexistence of both free and RECQL5-bound ECs in our sample. By selecting particles without RECQL5, we were able to solve the structure of a free EC containing only Pol II and the nucleic acid scaffold at 2.6 Å overall resolution (EC_Free_, **Fig. 1c**, **Table 1**, **Extended Data Figs. 1 and 2ab**). For the RECQL5-containing particles, we observed extensive flexibility in RECQL5, especially in its helicase domain (**Extended Data Figs. 3 and 4a,b**). This finding is in accordance with our previous observations, based on low-resolution negative-stain EM^22^, that RECQL5’s helicase domain can occupy a range of positions spanning up to a 60° arc around the downstream double-stranded DNA (dsDNA). To improve the resolution in the RECQL5 helicase domain and more clearly visualize its contacts with Pol II, we used a data processing approach in which 3D classification in a mask focused on the helicase domain is performed on signal-subtracted particles (detailed in **Methods**, **Extended Data Fig. 3**). For each conformation of the helicase domain identified in this manner, particle subtraction was reverted and the structure of the full complex was refined. This approach proved successful in addressing the conformational heterogeneity present in our dataset and yielded the structure of EC_REC-Apo_ at 3.2 Å overall resolution (**Fig. 1d**, **Table 1**, **Extended Data Fig. 2c**). The quality of our cryo-EM maps enabled us to model all domains of RECQL5 present (helicase domain, RQC domain, and IRI module), in addition to Pol II and the nucleic acid scaffold (**Fig. 1e,f**).

### RECQL5 contacts the Pol II EC through multiple domains

Our EC_REC-Apo_ structure shows how RECQL5 engages with the EC (**Fig. 1d,e**). There are multiple points of contact between RECQL5 and Pol II mediated by both the IRI module (both the αN helix and KIX domain) as well as the helicase D2 subdomain (**Fig. 1f**). We focused first on the IRI module interactions. To improve the resolution in this region, we reprocessed this dataset, classifying in the IRI module region instead of the helicase domain (**Extended Data Fig. 5**). This approach greatly improved the map’s local resolution at the IRI–Pol II interface, enabling us to trace the polypeptide chain (**Fig. 2b–d**, **Table 1**, **Extended Data Fig. 2d**). The KIX domain binds to the RPB1 lower jaw at the site predicted previously based on its homology to TFIIS^22,30^ and is anchored through interactions mediated by its α1 and α3 helices (**Fig. 2c**). Several residues in α3 (N595, K598, and R610) participate in hydrogen bonding or ionic interactions with residues in RPB1. Additionally, residues in the top half of α3 (V593, L596, L602) and in α1 (K548, H552) mediate nonpolar interactions with RPB1. R544, which is situated in a loop between α1 and α2, forms salt bridges with D1241 and D1242 in RPB1. Our structure further shows that the αN helix binds to a shallow nonpolar groove in the top face of the RPB1 lower jaw (**Fig. 2d**). Several residues (W504, Y508, M512, and R515) stick into this groove to interact with RPB1, while two others (F507 and Q511) interact with residues in the wall of the groove.

**Fig. 2.**
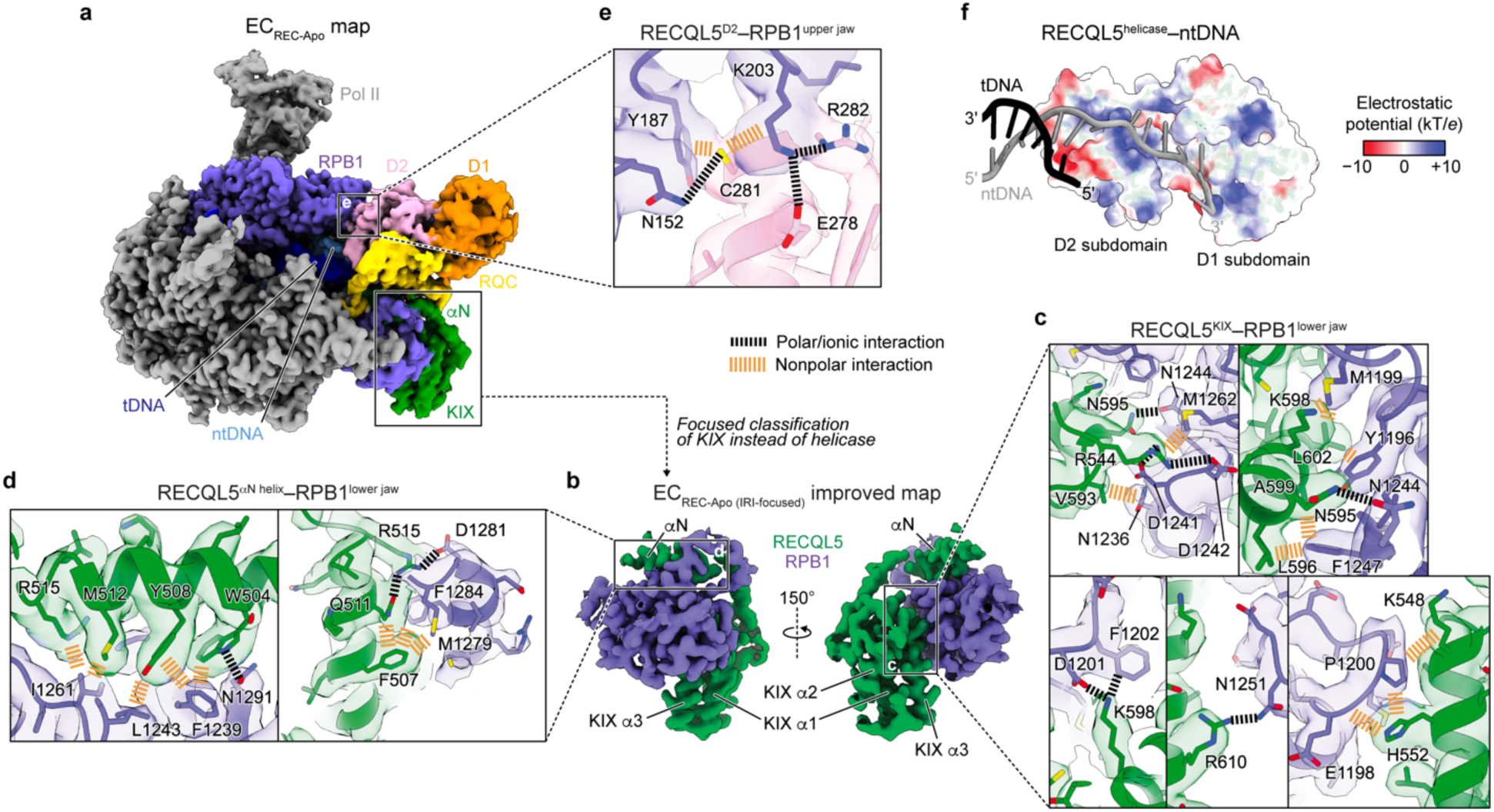
| RECQL5 contacts the Pol II EC through multiple domains. **a**. Full cryo-EM map for EC_REC-Apo_ highlighting the Pol II RPB1 subunit (purple) and the different domains in RECQL5 (colored as in Fig. 1a). **b**. Improved cryo-EM map (EC_REC-Apo (IRI-focused)_) of the RPB1 lower jaw region engaged with the RECQL5 IRI module (αN helix and KIX domain) (see **Extended Data Fig. 5** for details). **c–e.** Details of the interfaces between RECQL5 and Pol II in EC_REC-Apo_, with the cryo-EM map shown as a transparent surface and colored according to the fitted atomic model shown in ribbon, and with key residues shown in stick representation. Salt bridges and polar interactions are indicated in black, and nonpolar interactions are indicated in orange. Panels **c** and **d** show zoomed views of the map in **b** at the interfaces between the RPB1 lower jaw and the RECQL5 KIX domain (**c**), or the αN helix (**d**). Panel **e** shows a zoomed view of the map in **a** at the interface between the RPB1 upper jaw and the RECQL5 D2 helicase subdomain. **f.** View of the interface between the RECQL5 helicase domain and the single-stranded DNA. tDNA (black) and ntDNA (gray) are shown in cartoon representation, while the helicase domain is shown as a semi-transparent surface colored by electrostatic potential.

The isolated helicase domain of RECQL5 has been shown not to bind Pol II in an in vitro pulldown assay^22^. However, our present structure reveals a direct interaction between the RPB1 upper jaw and the D2 helicase subdomain in the context of the EC (**Fig. 2a**). This interface is stabilized by interactions between RECQL5 E278 and R282 and RPB1 K203, as well as by both polar and nonpolar interactions between RECQL5 C281 and RPB1 N152, Y187, and K203 (**Fig. 2e**). We note that the helicase conformation we describe in the EC_REC-Apo_ structure (stabilized in part by D2–Pol II interactions), obtained after rounds of local classification, likely represents the most stable of the many possible poses that it can adopt. Finally, the EC_REC-Apo_ structure shows that both helicase D1 and D2 subdomains of RECQL5 bind to the single-stranded ntDNA extension (**Fig. 2f**) via a positively charged channel. Overall, our structural analysis reveals the details of how RECQL5 contacts the Pol II EC.

### Free and RECQL5-bound Pol II exist in translocation intermediate states

Given that RECQL5 contacts both Pol II and the extruding ntDNA, we considered the possibility that it might affect the Pol II translocation state in the EC. Therefore, we examined the Pol II active site in the EC_Free_ versus EC_REC-Apo_ complexes. The good local resolution (2.3–3.1 Å) attained for the Pol II core region allowed us to assign unambiguously the DNA/RNA register (**Fig. 3a,b**). In EC_Free_, the tDNA *i* site base G14 is positioned a distance of 8.7 Å from the RPB1 bridge helix (Cα of RPB1 T854 taken as a reference point) (**Fig. 3c,d**). This distance is extremely close to the 8.6 Å distance observed for Pol II bound to the transcription inhibitor α-amanitin, which adopts a translocation intermediate conformation^29^ (PDB 2VUM), and is significantly different from the 6 Å distance observed for Pol II in a pre-translocation state^31^ (PDB 1I6H) (**Fig. 3e**). Moreover, when superimposed on the bridge helix, EC_Free_ showed great similarity to the Pol II–α-amanitin structure (RMSD = 0.5 Å) (**Extended Data Fig. 6**). These observations indicate that in the absence of RECQL5, Pol II adopts an intermediate conformation between the pre– and post-translocation states on our nucleic acid scaffold (**Fig. 3f**).

**Fig. 3.**
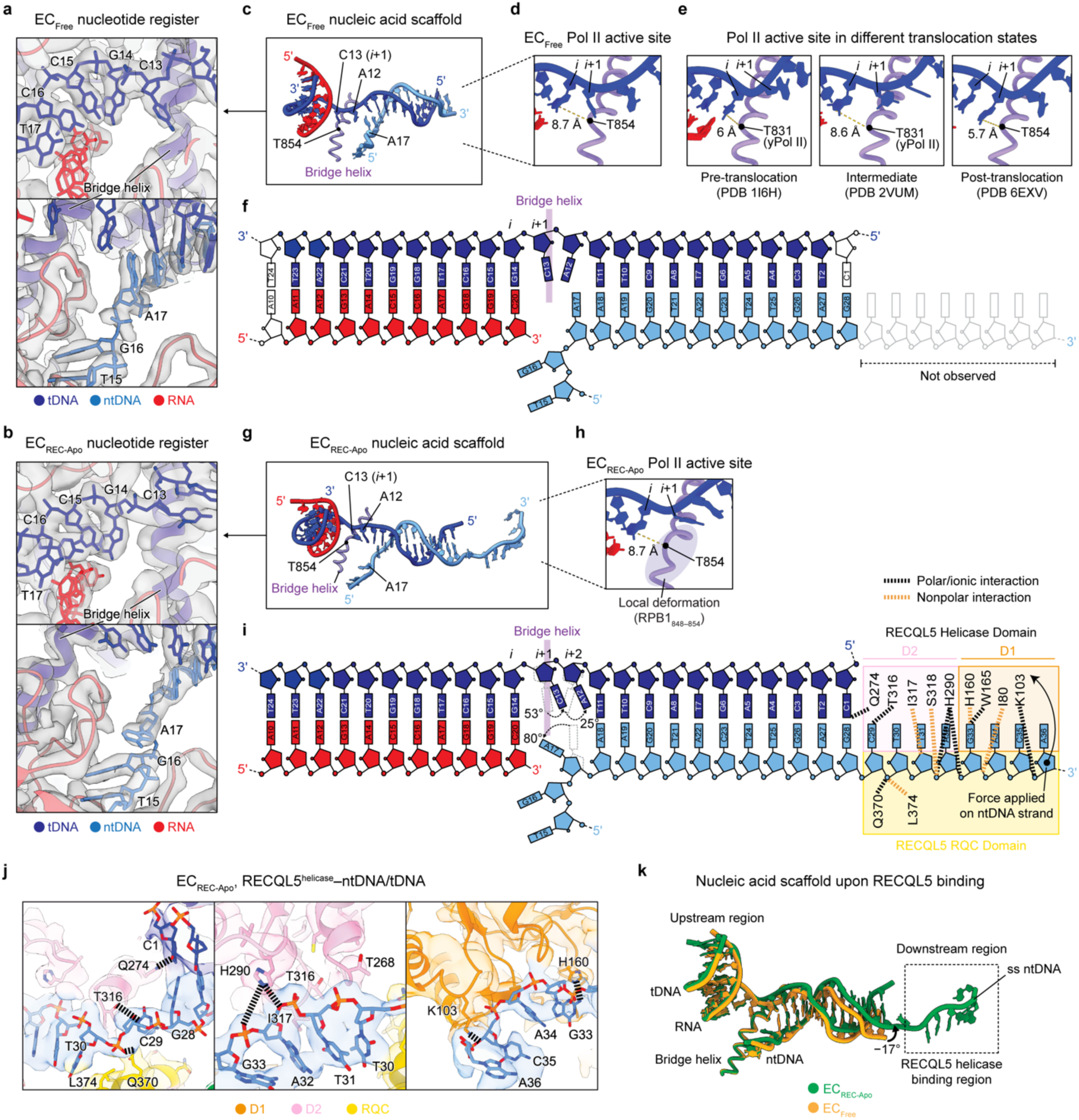
| Free and RECQL5-bound Pol II exist in translocation intermediate states. **a,b**. Regions near the Pol II active site upstream of the bridge helix (top) and downstream of the bridge helix (bottom) in the EC_Free_ (**a**) and EC_REC-Apo_ structures (**b**). Cryo-EM density is shown as a transparent gray surface, with the fitted atomic model in ribbon representation and the DNA bases in stick representation. Key nucleotides are labeled. Colors are indicated. **c**. Model of the nucleic acid scaffold and Pol II bridge helix in the EC_Free_ structure. Important nucleotides are highlighted. **d**. Close up of the EC_Free_ Pol II active site shown in **c** highlighting the distance between the *i* site nucleotide base and the RPB1 T854 residue in the bridge helix. **e**. Corresponding views of the Pol II active site in ECs where Pol II is in a pre-translocation conformation (PDB 1I6H, left), translocation intermediate conformation (PDB 2VUM, middle), or post-translocation conformation (PDB 6EXV, right). The distance from the active site nucleotide (*i* or *i*+1) and the RPB1 T854 residue (T831 in yeast RPB1) is indicated in each case. **f**. Schematic depicting the nucleic acid scaffold in EC_Free_ highlighting the nucleotides that are resolved in the structure and the position of the bridge helix. **g**. Model of the nucleic acid scaffold and Pol II bridge helix in EC_REC-Apo_. Important nucleotides are highlighted. **h**. Close up of the EC_REC-Apo_ Pol II active site shown in **g** highlighting the distance between the *i* site nucleotide base and the RPB1 T854 residue in the bridge helix. **i**. Schematic depicting the nucleic acid scaffold in EC_REC-Apo_ highlighting the nucleotides that are resolved in the structure, the position of the bridge helix, and the residues in the RECQL5 helicase domain that interact with the nucleic acids. Salt bridges and polar interactions are indicated in black, and nonpolar interactions are indicated in orange. **j**. Views of the downstream region in EC_REC-Apo_ highlighting molecular interactions between the RECQL5 helicase domain and the ntDNA. The cryo-EM map is shown as a transparent surface, with the fitted atomic model in ribbon representation and the residue side chains and DNA bases in stick representation. Salt bridges and polar interactions are indicated with black dashed lines. Residues of interest are explicitly labeled. **k**. Superimposition of the nucleic acid scaffolds from the EC_Free_ (yellow) and EC_REC-Apo_ (green) structures. The EC_REC-Apo_ structure shows a −17° rotation corresponding to partial DNA unwinding compared to the EC_Free_ structure.

Compared to EC_Free_, EC_REC-Apo_ displays a slightly reorganized Pol II active site (**Fig. 3g**). In EC_REC-Apo_, the *i* site tDNA base G14 appears 8.7 Å away from the bridge helix, similar to its positioning in EC_Free_ (**Fig. 3h**). However, we observe a local distortion of the bridge helix in EC_REC-Apo_, in contrast to in the EC_Free_ and Pol II–α-amanitin structures (**Fig. 3h**, **Extended Data Fig. 6**). Moreover, while we did not observe density in EC_Free_ corresponding to the single-stranded ntDNA (indicating it is not stably tethered to Pol II), in EC_REC-Apo_ this region is stabilized through contacts with the RECQL5 helicase and RQC domains (**Figs. 2f and 3i,j**). Within the helicase domain, most interactions with the ssDNA are mediated by residues in the D2 subdomain, with fewer interactions contributed by the D1 subdomain. In this interaction mode, the RECQL5 helicase appears to adopt a ‘pushing and unwinding’ state where it pushes the ssDNA and concomitantly bends the downstream dsDNA end by −17°, leading to its partial unwinding (**Fig. 3k**). This local DNA bending causes distortions of the B-DNA helix structure that are propagated upstream until reaching the DNA/RNA hybrid region, which itself does not display structural rearrangements. Interestingly, when inspecting the Pol II active site in the EC_REC-Apo_ structure, the tDNA bases C13 (*i+1* site) and A12 (*i+2* site) appear significantly rotated towards the downstream region, by 53° and 25° relative to their positioning in the EC_Free_, respectively (**Fig. 3i**). Additionally, the non-template DNA base A17, located immediately downstream to the bridge helix, appears rotated 80° towards the upstream region, relative to its position in EC_Free_. Altogether, these observations indicate that Pol II in the EC_REC-Apo_ structure also adopts an intermediate conformation similar to that observed in EC_Free_, albeit with a slightly altered active site configuration.

### Structures of ECs with nucleotide-bound RECQL5 helicase

In light of these observations, we considered whether the Pol II translocation state might change depending on the nucleotide-binding state of the RECQL5 helicase domain. Previous studies have shown that RECQL5’s helicase activity is not necessary to inhibit Pol II transcription in vitro^21,22^. However, structural work has shown that the helicase domain changes conformation upon ADP binding, with the D1 subdomain rotating ∼20° relative to the D2 subdomain^32^. Those structures were obtained in the absence of DNA, and therefore it remains unclear whether this conformational change can take place in the context of an EC. In turn, we wondered if a helicase conformational change could affect the translocation state of Pol II.

To investigate these questions, we assembled and purified RECQL5 ECs via a pulldown approach, using wild-type RECQL5 in the presence of either adenylyl-imidodiphosphate (a non-hydrolyzable ATP analog also known as AMPPNP) or ADP (**Extended Data Fig. 7a,b**). We collected cryo-EM data for these complexes and used a similar processing workflow as for EC_REC-Apo_ to solve their structures, which we refer to as EC_REC-AMPPNP_ and EC_REC-ADP_ (**Table 1**, **Extended Data Figs. 8a–c, 2e,f, 9, and 10**). Comparison of these structures with the EC_REC-Apo_ structure showed that the IRI modules were almost identical in conformation (**Extended Data Fig. 8d**), with root-mean-squared deviations (RMSDs) less than 1 Å (0.554 Å for EC_REC-Apo_ versus EC_REC-AMPPNP_, 0.852 Å for EC_REC-Apo_ versus EC_REC-ADP_). These observations support the notion that the IRI module serves to anchor RECQL5 to Pol II throughout the helicase domain’s nucleotide binding and hydrolysis cycle.

### Helicase nucleotide binding induces a post-translocation Pol II conformation

AMPPNP binding induced noticeable conformational changes within both the Pol II active site and the helicase–ntDNA interaction region. As with the previous structures, we were able to confidently assign the DNA/RNA register in EC_REC-AMPPNP_ and therefore identify the locations of the *i* and *i*+1 nucleotides in the Pol II active site (**Fig. 4a–c**). We observed several major conformational changes in EC_REC-AMPPNP_ compared to EC_REC-Apo_. First, in the tDNA strand, the A12 base (*i+2* site) appears rotated by 16° in the upstream direction, while the ntDNA base A17 is rotated by 92° in the downstream direction (**Fig. 4d**). Strikingly, the tDNA base C13 (*i+1* site) displayed a large 93° rotation towards the upstream region to be located immediately upstream of the bridge helix, positioned 5.4 Å away from it (**Fig. 4c**). Comparing this EC_REC-AMPPNP_ structure (**Fig. 4c**) to previous ECs (**Fig. 3e**), we conclude that Pol II adopts a post-translocation conformation in this complex.

**Fig. 4.**
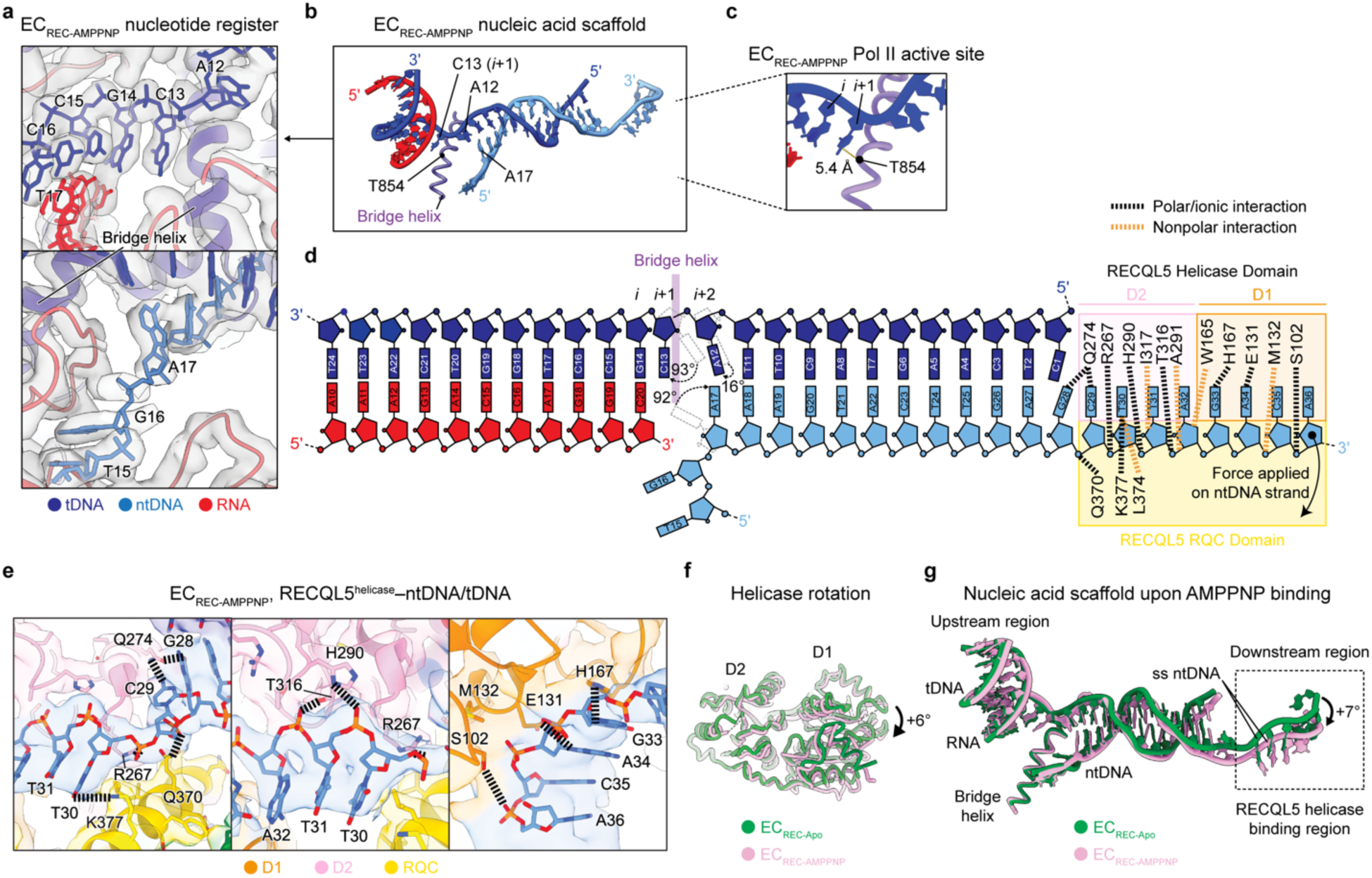
| Helicase nucleotide binding induces a post-translocation Pol II conformation. **a**. Regions near the Pol II active site upstream of the bridge helix (top) and downstream of the bridge helix (bottom) in EC_REC-AMPPNP_. The full cryo-EM map for EC_REC-AMPPNP_ is shown in **Extended Data Fig. 8a**. Cryo-EM density is shown as a transparent gray surface, with the fitted atomic model in ribbon representation and the DNA bases in stick representation. Key nucleotides are labeled. Colors are indicated. **b**. Model of the nucleic acid scaffold and Pol II bridge helix in the EC_REC-AMPPNP_ structure. Important nucleotides are highlighted. **c**. Close up of the EC_REC-AMPPNP_ Pol II active site shown in **b** highlighting the distance between the *i*+1 site nucleotide base and the RPB1 T854 residue in the bridge helix. **d**. Schematic depicting the nucleic acid scaffold in EC_REC-AMPPNP_ highlighting the nucleotides that are resolved in the structure, the position of the bridge helix, and the residues in the RECQL5 helicase domain that interact with the nucleic acids. Salt bridges and polar interactions are indicated in black, and nonpolar interactions are indicated in orange. **e**. Views of the downstream region in EC_REC-AMPPNP_ highlighting molecular interactions between the RECQL5 helicase domain and the ntDNA. The cryo-EM map is shown as a transparent surface, with the fitted atomic model in ribbon representation and the residue side chains and DNA bases in stick representation. Salt bridges and polar interactions are indicated with black dashed lines. Residues of interest are explicitly labeled. **f**. Comparison of the RECQL5 helicase conformation in EC_REC-Apo_ (green) versus EC_REC-AMPPNP_ (pink). The two structures are aligned on the helicase D2 subdomain, revealing a +6° relative rotation of the D1 subdomain. **g**. Superimposition of the nucleic acid scaffolds from the EC_REC-Apo_ (green) and EC_REC-AMPPNP_ (pink) structures. The EC_REC-AMPPNP_ structure shows a +7° rotation corresponding to DNA rewinding compared to the EC_REC-Apo_ structure.

To understand how the post-translocation conformation is promoted by AMPPNP binding, we examined the RECQL5 helicase domain’s interactions with the single-stranded ntDNA. In EC_REC-AMPPNP_, the single-stranded ntDNA is contacted by additional residues compared to EC_REC-Apo_ (**Figs. 3h and 4d,e**). Moreover, the helicase D1 subdomain is rotated +6° around the D2 subdomain relative to EC_REC-Apo_ (**Fig. 4f**), resulting in a new interaction mode that we refer as a ‘pulling and rewinding’ state. Here, the RECQL5 helicase appears to pull the single-stranded ntDNA around its 3’ end, concomitantly bending the dsDNA downstream region by +7° and leading to its partial rewinding (**Fig. 4g**).

We also examined the effects of RECQL5 binding to ADP. The cryo-EM structure of EC_REC-ADP_ showed an overall architecture similar to EC_REC-Apo_ and EC_REC-AMPPNP_ (**Figs. 1d,e** **and Extended Data Fig. 8a,b**). In EC_REC-ADP_, neither the αN helix nor the KIX domain show major changes relative to their positions in EC_RECApo_ or EC_REC-AMPPNP_ (**Extended Data Fig. 8d**). The main difference with respect to the two other states analyzed was that the RECQL5 helicase domain in EC_REC-ADP_ exhibits high orientational flexibility relative to Pol II, which led to lower local resolution of this region (7–8.4 Å; **Extended Data Fig. 2f**), despite extensive data analysis (**Extended Data Fig. 10**). Moreover, the density assigned to the D1 subdomain remained blurrier than that for the D2 subdomain (**Extended Data Fig. 8b**), allowing only partial modelling of D1 and the single-stranded ntDNA it interacts with. These results suggest that in the ADP state, the RECQL5 helicase D1 and D2 subdomains may be more dynamic. Notably, we did not observe the large rotation between these domains seen in crystallographic structures of the free RECQL5 helicase domain^32^. We posit that this may be due to constraints imposed by DNA– and Pol II-binding within the context of the EC. Inspection of the Pol II active site in EC_REC-ADP_ shows that Pol II is in the post-translocation state (**Extended Data Fig. 8e,f**), as we observed in EC_REC-AMPPNP_ (**Fig. 4c**). This result suggests that ATP hydrolysis and phosphate release by RECQL5 do not provide the energy associated with changes in the Pol II translocation configuration, but rather the conformational change of RECQL5 upon ATP binding likely does.

## Discussion

RECQL5 is a regulator of transcription elongation known to interact directly with Pol II, thereby playing important roles in suppressing genome instability. Nevertheless, detailed structural and mechanistic insights into its molecular interplay with Pol II are currently lacking. In this study, we solved several cryo-EM structures of Pol II ECs complexed with RECQL5 in different nucleotide-binding states of the helicase. Our structures reveal the molecular interactions that stabilize RECQL5 binding to the Pol II EC. The RECQL5 IRI module, encompassing the αN helix and KIX domain, wraps around the surface of the RPB1 lower jaw with exquisite molecular complementarity, serving as an anchor point of RECQL5 on Pol II. Meanwhile, the helicase domain binds to the RPB1 upper jaw and downstream DNA. Below, we discuss the potential implications of our work within the context of the distinct biological roles that have been ascribed to RECQL5 in the cell.

Our structural results suggest that the RECQL5 helicase domain may exert its regulatory effects on transcription via two distinct mechanisms. First, the helicase domain binds to DNA downstream of the transcribing Pol II, thus likely acting as a steric roadblock that slows down transcription, as previously proposed^22^ (**Fig. 5a**). The resolution of our cryo-EM structures allowed us to map critical molecular interactions of the RECQL5 helicase domain with the single-stranded DNA and with the upper jaw of the Pol II. We also observed extensive conformational flexibility for the helicase domain in all of our datasets, most pronounced for EC_REC-ADP_, while the position of the IRI module remained unaltered between the EC_REC-Apo_, EC_REC-AMPPNP_, and EC_REC-ADP_ structures. This result indicates that the IRI–RPB1^lower^ ^jaw^ interaction is stable and anchors the flexible and more loosely engaged RECQL5 helicase domain to the Pol II EC. This role may be bolstered by the interaction between the RECQL5 SRI domain (not included in our truncated RECQL5 construct) and the disordered Pol II C-terminal domain, which may help increase the local concentration of RECQL5 around elongating Pol II^19^. Overall, a helicase roadblock mechanism agrees with previous studies showing that RECQL5 can inhibit transcription in vitro^21,22^ as well as globally slow down elongation rates in cells^23^.

**Fig. 5.**
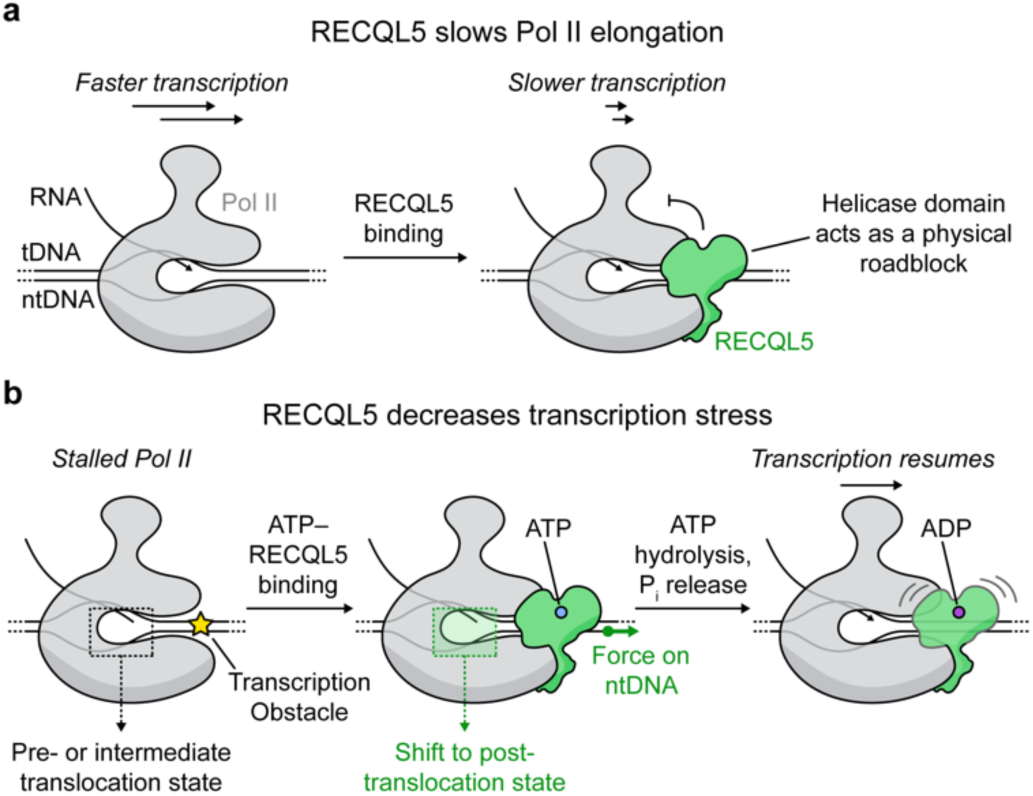
| Proposed mechanisms of transcription regulation by RECQL5. **a**. Schematic depicting the function of RECQL5 as a negative elongation factor. RECQL5 globally slows down Pol II transcription, presumably by binding to downstream DNA and acting as a roadblock for Pol II advancement. **b.** Schematic of a second proposed function of the RECQL5 helicase domain. During Pol II stalling, RECQL5 may help to reset the translocation status of Pol II through its interactions with the downstream DNA. Initial binding of ATP–RECQL5 to the EC is accompanied by a pulling force on the ntDNA. This mechanical stress is propagated along the DNA to the Pol II active site, shifting it to a post-translocation state that is ready to incorporate a new incoming nucleotide into the growing RNA. Subsequent ATP hydrolysis and phosphate (P_i_) release by the RECQL5 helicase domain increases its conformational flexibility, which may facilitate transcription restart and bypassing of the helicase roadblock. We propose that this mechanism may help explain how RECQL5 decreases transcription stress in cells.

Second, our findings show that RECQL5 has the ability to modulate Pol II’s translocation status, indicating that it could help to restart stalled ECs. In EC_REC-APO_, the RECQL5 helicase domain appears to push the downstream DNA against Pol II, bending it by −17° and partially unwinding it. Upon AMPPNP binding in EC_REC-AMPPNP_, the RECQL5 helicase domain undergoes an internal conformational change (D2 subdomain rotates +6°) that reorganizes its contacts with the single-stranded ntDNA and pulls it away from Pol II, partially rewinding the DNA. These observations are consistent with the known property of double-stranded DNA to exhibit negative coupling between twisting and stretching under small distortions, overwinding when stretched and underwinding when contracted^33^. This twisting-stretching inverse coupling is of particular importance for DNA-binding proteins that exploit this property by causing local distortions of the B-DNA geometry upon binding^33^. Our structures show that these pushing and pulling actions induced by RECQL5 in different nucleotide-binding states apply torques on the DNA that propagate upstream to the Pol II active site. Crucially, in the case of AMPPNP-bound RECQL5, this mechanism of action induces Pol II to adopt a post-translocation state.

Notably, a previous study showed that RECQL5 not only slows down transcription elongation but also reduces transcription stress, which is linked to genome instability^23^. Based on our structures, we propose that the RECQL5 helicase domain may act mechanically on a Pol II that has been stalled or paused during transcription (**Fig. 5b**). This process begins when RECQL5, complexed to ATP, binds the Pol II EC. The helicase domain engages with the RPB1 upper jaw and downstream DNA, pulling outwards on the latter, generating torsion on the DNA that is allosterically transmitted towards the Pol II active site. There, it stabilizes the post-translocation conformation, which becomes ready to incorporate a new incoming nucleotide into the growing RNA. This model resolves the apparent contradiction of how a factor that acts as a roadblock can nonetheless favor transcription elongation by lowering the levels of Pol II stalling and backtracking. ATP hydrolysis and phosphate release by the helicase domain do not alter the Pol II translocation state. Instead, they serve to destabilize the binding of the helicase domain to the EC, which becomes more flexible. We speculate that this loose binding between the helicase subdomains and the EC, in turn, facilitates Pol II’s forward progress. We propose that the effect is to facilitate the restarting of transcription in the event of Pol II pausing, thereby decreasing transcription stress and associated genome instability (**Fig. 5b**).

On the other hand, more recent evidence has emerged that RECQL5 plays a critical role in the resolution of transcription-replication conflicts. Specifically, RECQL5 helicase activity is necessary to disassemble RAD51 filaments from the stalled replication fork, an important step in replication restart^13–15^. An important outstanding question is whether and how RECQL5’s role in transcriptional regulation is connected to its function in resolving stalled replication forks. In this context, it is tempting to speculate that the anchoring of RECQL5 to elongating Pol II ECs—via its IRI module and SRI domain—may serve as a means to localize it to transcription-replication conflicts, thereby recruiting it for RAD51 filament disassembly. Additionally, RECQL5’s proposed ability to help restart stalled Pol II transcription may also contribute to the overall process of resolving such collisions. Understanding the molecular basis for RECQL5’s function in resolving transcription-replication conflicts will be an important goal for future studies. Moreover, although RECQL5’s unstructured sequence is known to contain a RAD51-interacting region^11^, structural insights are lacking into both this interaction and the process of filament disassembly. These future studies will further elucidate RECQL5’s disparate genome-protective roles.

## Author Contributions

E.N., A.J.F.A., and N.Z.L. conceived the study and designed experiments. A.J.F.A. and N.Z.L. prepared cryo-EM samples and collected and analyzed cryo-EM data. A.J.F.A. developed the cryo-EM processing approach. N.Z.L., J.F., and S.A.K. purified proteins. P.G. assisted with graphene oxide grid fabrication and provided technical advice regarding cryo-EM sample preparation. S.A.K. performed initial sample preparation, and B.K. prepared cryo-EM grids and collected the initial cryo-EM dataset. E.N. supervised the study. A.J.F.A., N.Z.L., and E.N. wrote the manuscript, and all authors edited the manuscript.

## Acknowledgments

We thank the members of the Nogales Lab for important discussions; D. Toso and R. Thakkar at the Cal-Cryo EM facility at QB3-Berkeley for help with EM data acquisition; F. Burgos-Bravos for biochemical advice; P. Tobias, K. Stine, and V. Marquez for computing support; and the UC Berkeley Cell Culture Facility for providing HeLa cells. We thank C. Bustamante for his feedback on the manuscript. This work was supported by the National Institutes of Health (NIH) grant R35-GM127018 to E.N. E.N. is a Howard Hughes Medical Institute (HHMI) Investigator.

## Extended Data Figures

**Extended Data Fig. 1.**
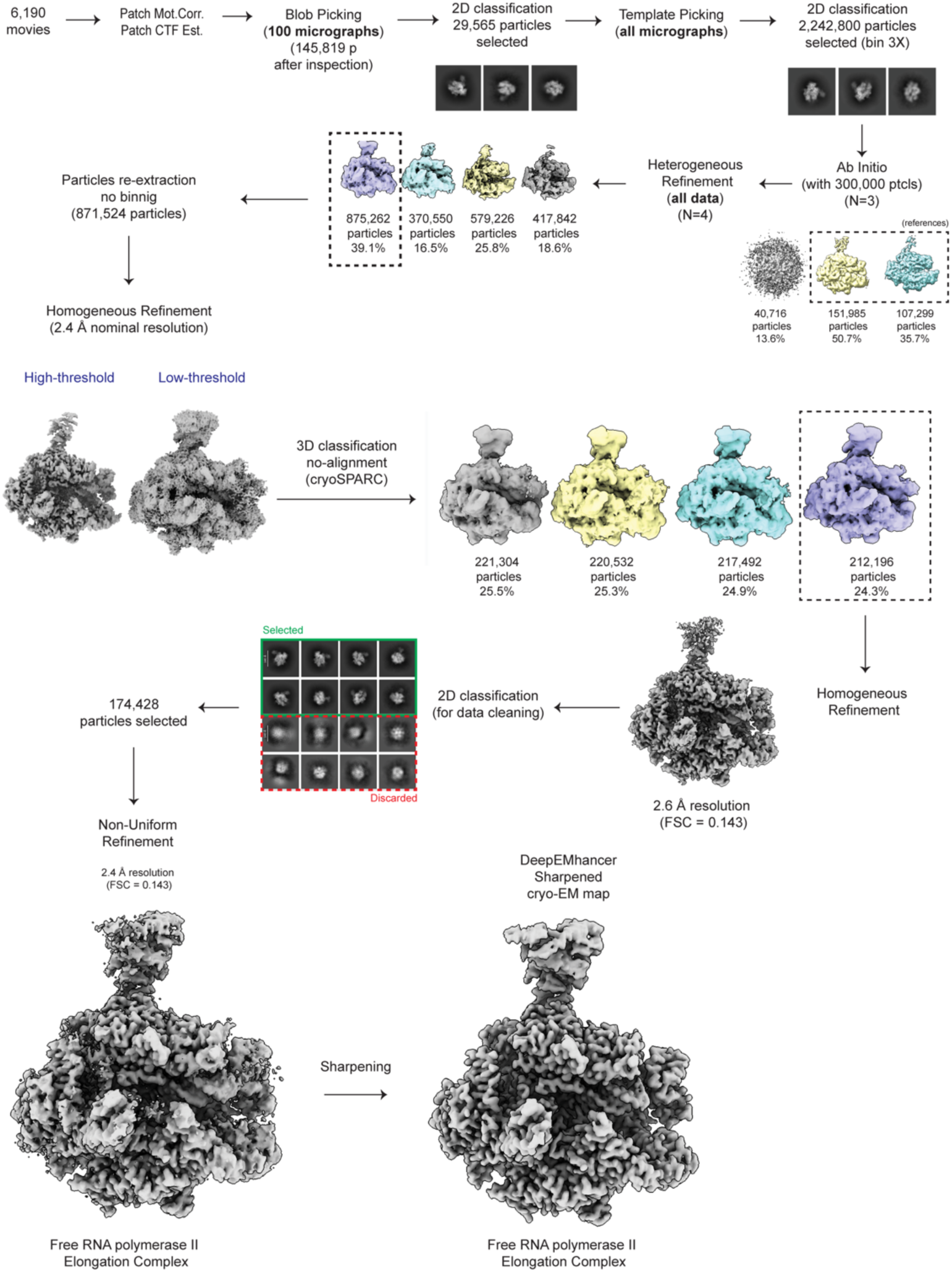
| Cryo-EM processing workflow for EC_Free_.

**Extended Data Fig. 2.**
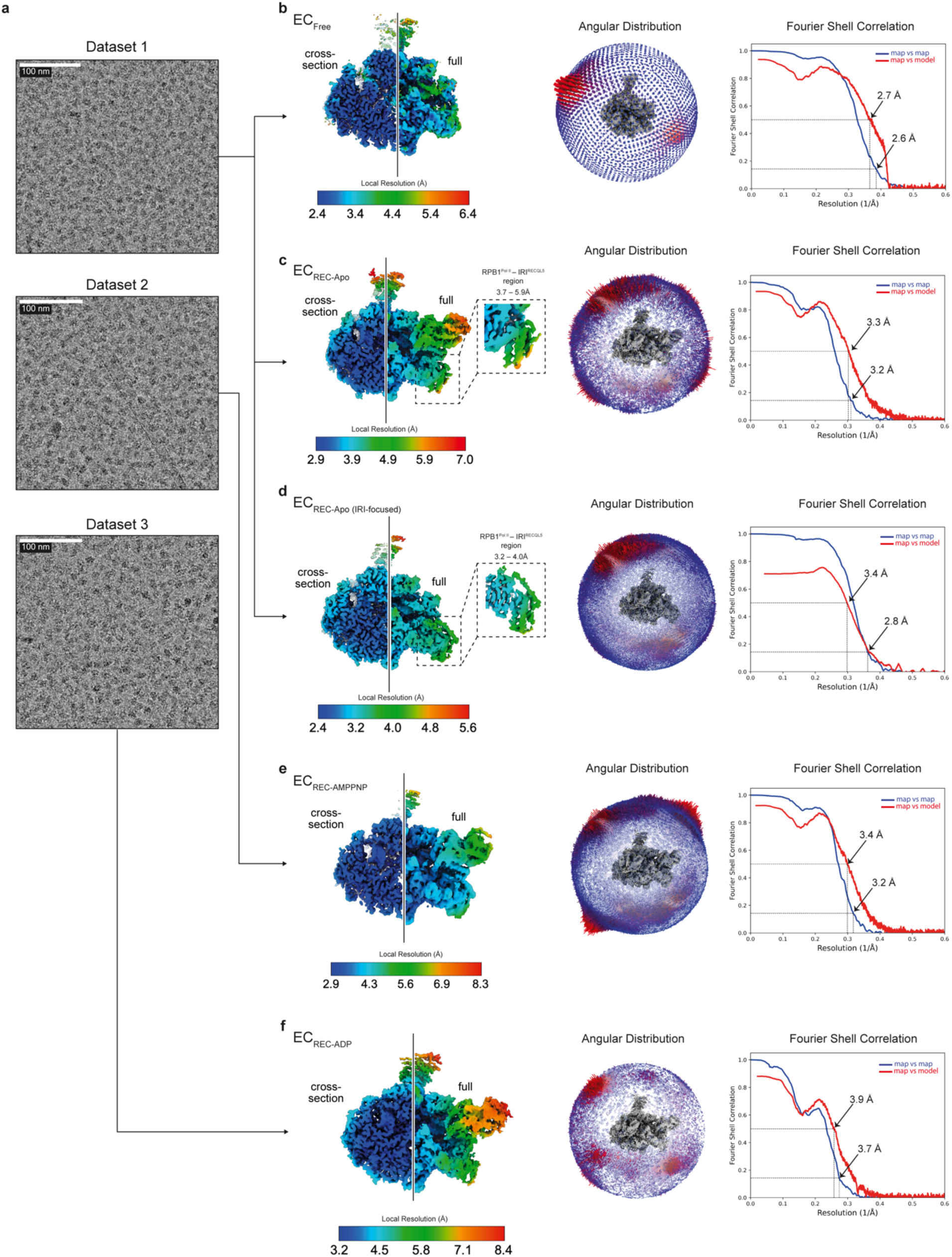
| Cryo-EM data collection and resolution estimation. **a**. Examples of cryo-EM corrected micrographs for the data sets analyzed. Scale bars are indicated. **b–f.** Unsharpened cryo-EM map colored by local resolution as indicated in the color key (left), angular distribution plot (middle), and Fourier Shell Correlation (FSC) map vs map and map vs model plots (right) for the EC_Free_ (**b**), EC_REC-Apo_ (**c**), EC_REC-Apo (IRI-focused)_ (**d**), EC_REC-AMPPNP_ (**e**), and EC_REC-ADP_ (**f**) structures. For the local resolution plots, the line separates a cross-section on the left and a view of the cryo-EM map surface on the right. In **c** and **d**, the insets show a close-up view of the RPB1 lower jaw and IRI module colored by local resolution.

**Extended Data Fig. 3.**
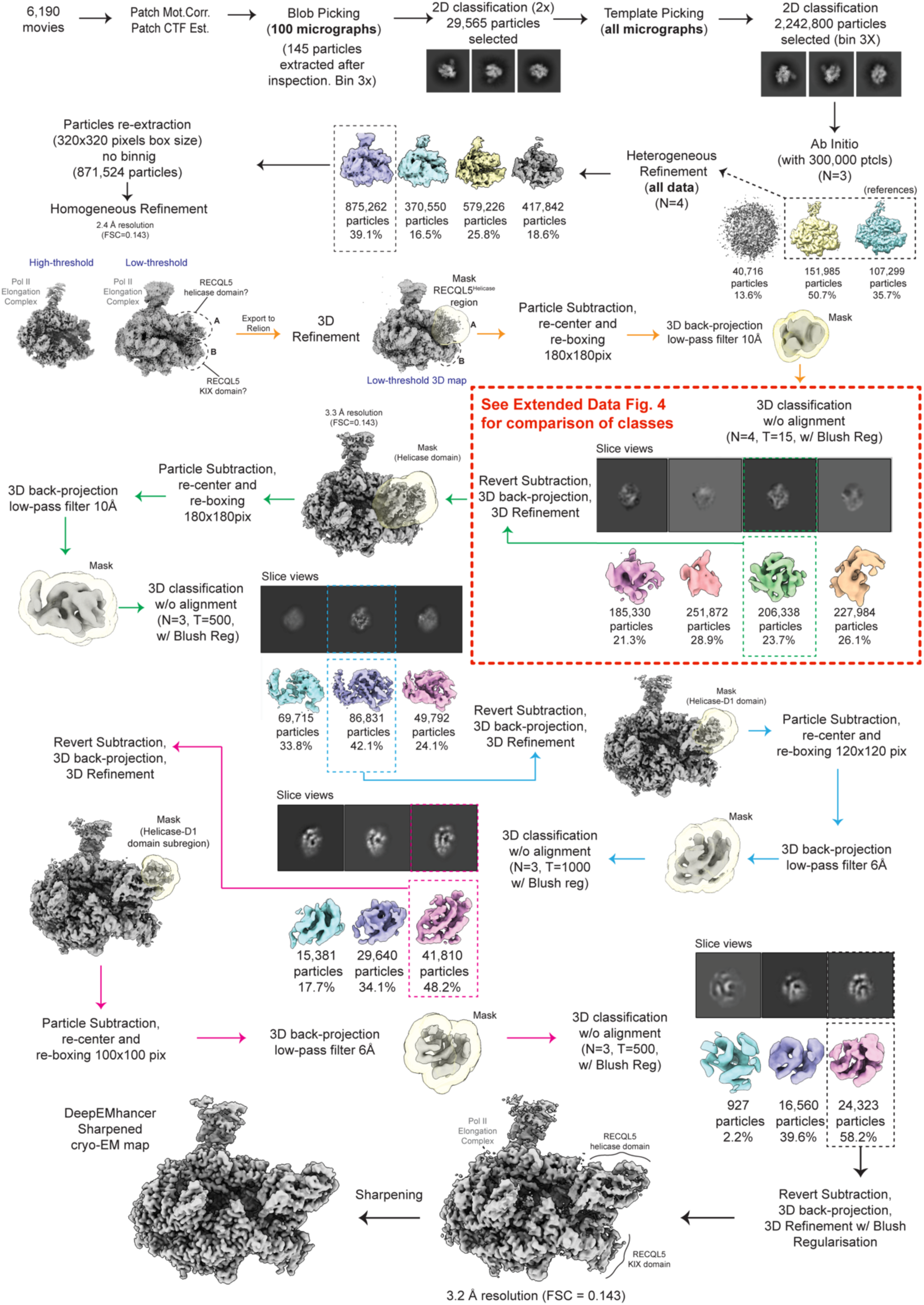
| Cryo-EM processing workflow for EC_REC-Apo_.

**Extended Data Fig. 4.**
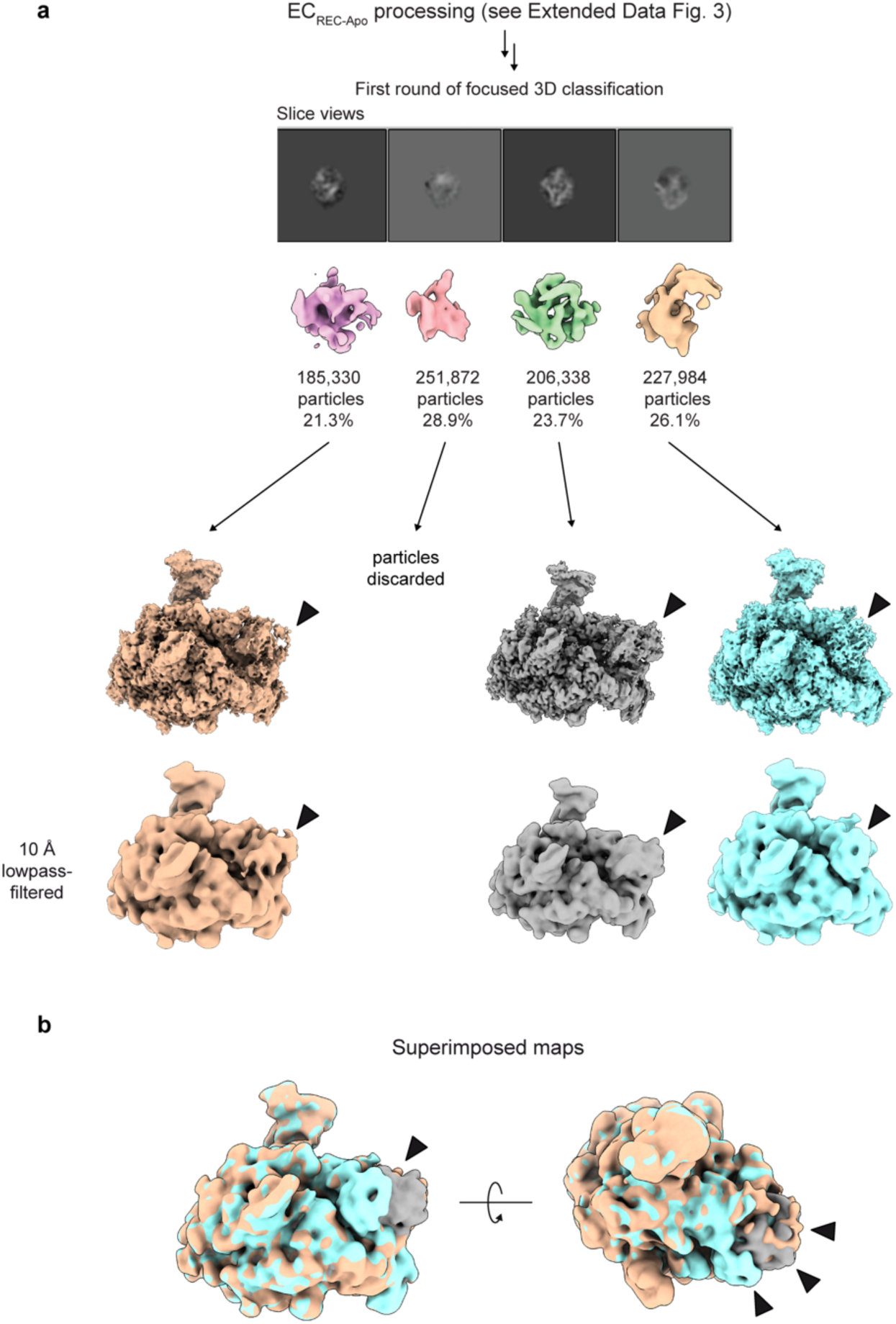
| RECQL5 helicase domain is flexible in the EC_REC-Apo_ dataset. **a**. Results from the first focused 3D classification step (see **Extended Data Fig. 3**). Classification was carried out on subtracted particles, focusing on the helicase domain. After classification, subtraction was reverted and particles were back-projected to obtain 3D reconstructions, which were subsequently refined. In three classes showing strong signal for the helicase domain, which are depicted in orange, gray, and cyan, the helicase domain occupies different positions at the Pol II DNA entry site. The gray map represents the class that was selected for further classification and refinement to obtain the final EC_REC-Apo_ structure. Arrows denote density corresponding to the helicase domain. **b.** Superimposition of the three lowpass-filtered maps in **a** with arrows highlighting the different positions the helicase domain adopts.

**Extended Data Fig. 5.**
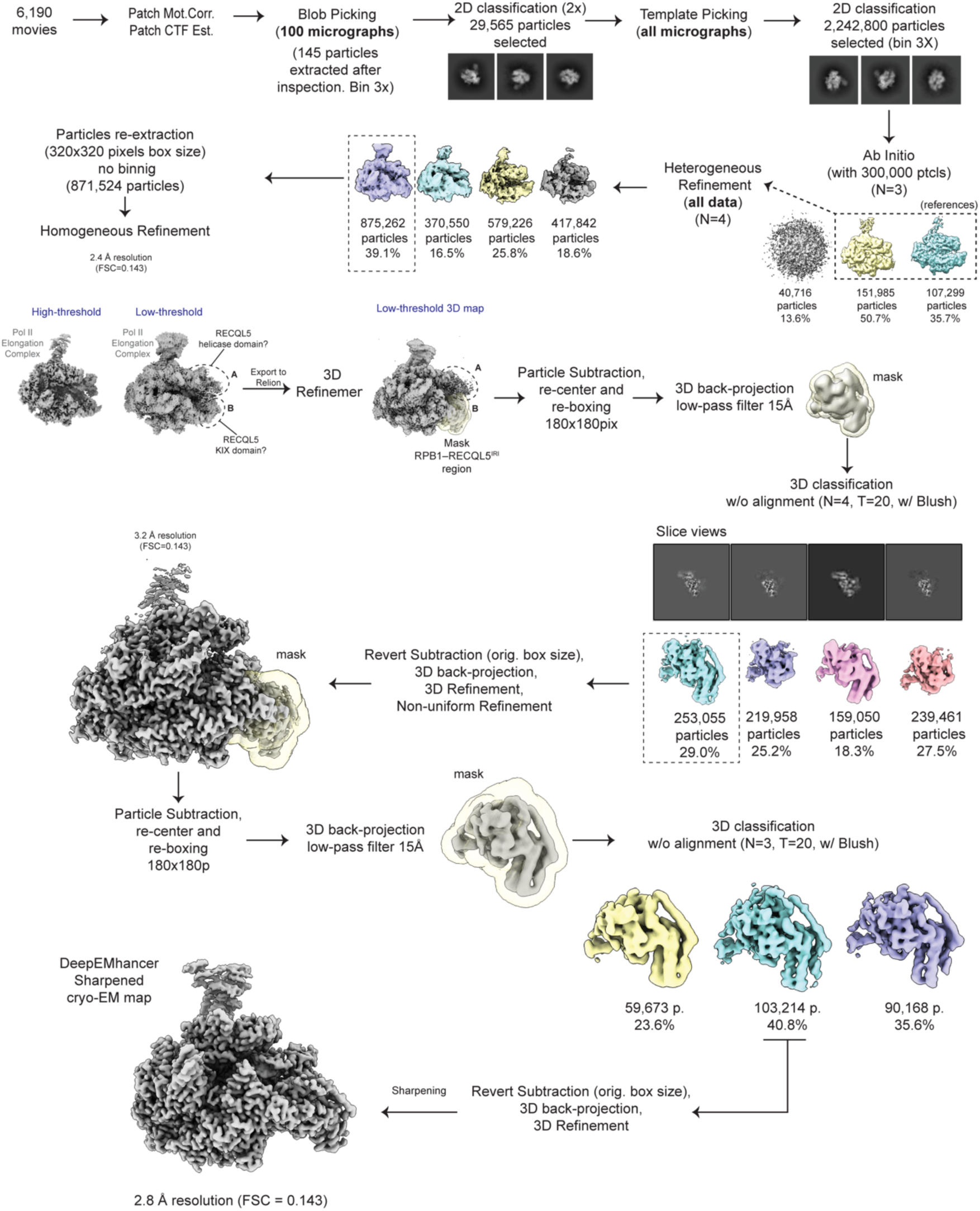
| Cryo-EM processing workflow for EC_REC-Apo (IRI-focused)_.

**Extended Data Fig. 6.**
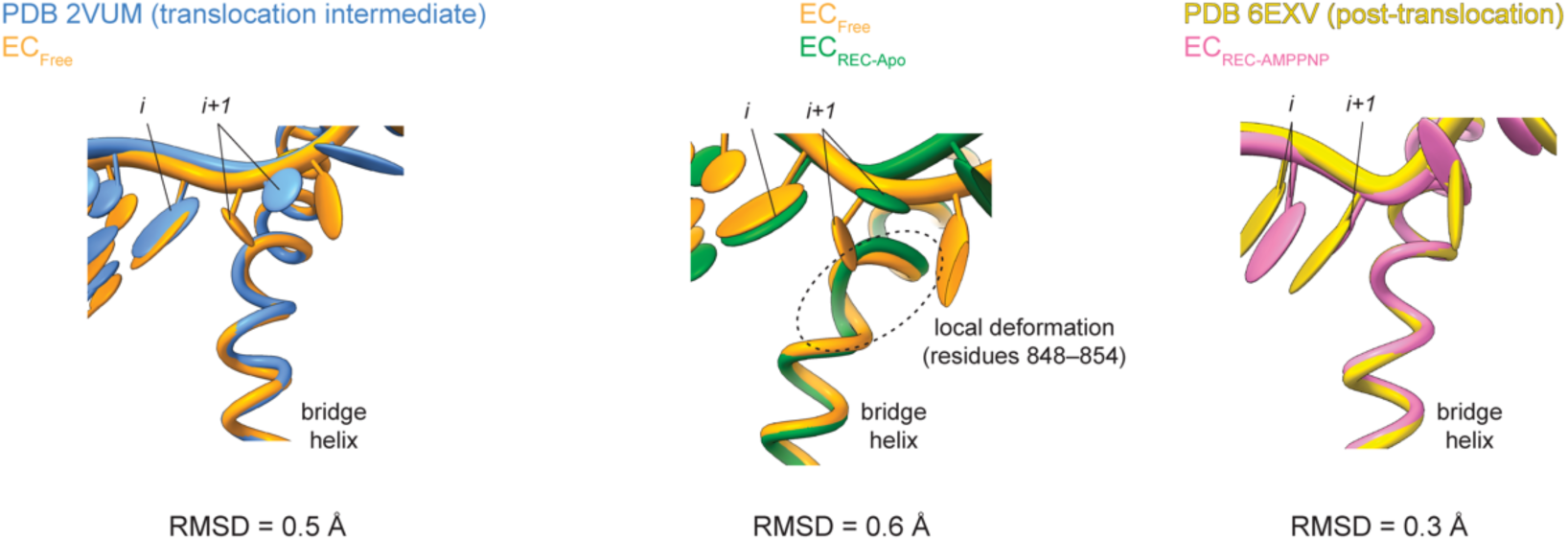
| Comparison of Pol II elongation complexes. Superposition of the bridge helix in EC_Free_ vs PDB_2VUM_ (left), EC_Free_ vs EC_REC-Apo_ (middle), and EC_REC-AMPPNP_ vs PDB_6EXV_ (right). RMSDs between the structures for each comparison are shown below. Visual inspection of the tDNA base in the *i and i+1* sites across the models (**Figs. 3d,e,h and 4c**), as well as the RMSDs reported here, indicate that EC_Free_ and EC_REC-Apo_ adopt distinct translocation intermediates, while EC_REC-AMPPNP_ adopts a post-translocation state. Color code is indicated.

**Extended Data Fig. 7.**
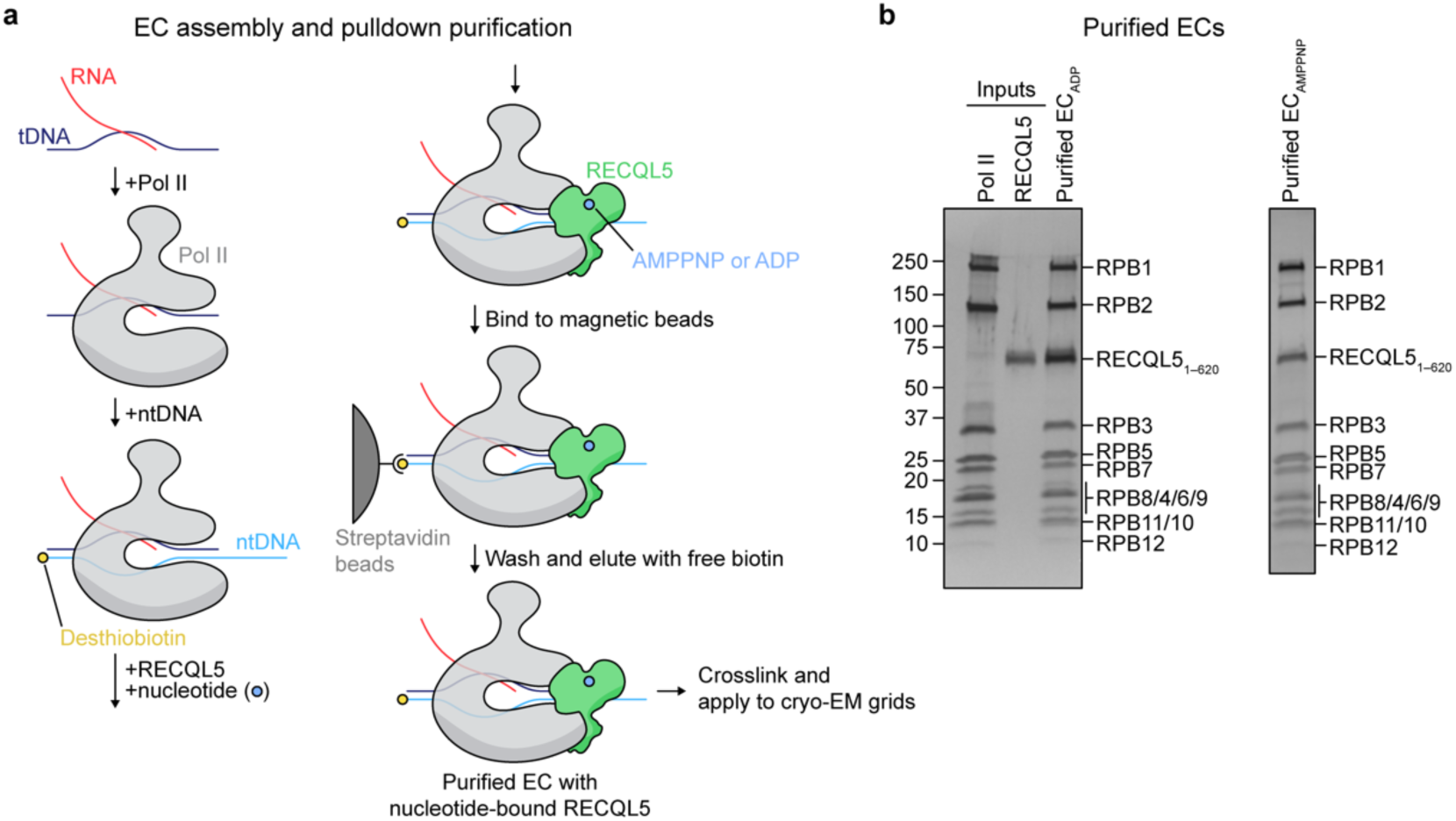
| Assembly and purification of EC_REC-AMPPNP_ and EC_ADP_ complexes. **a**. Schematic depicting pulldown strategy to reconstitute and purify Pol II ECs with nucleotide-bound RECQL5. **b**. SDS-PAGE analysis of pulldown inputs and the final purified EC_REC-ADP_ (left) and EC_REC-AMPPNP_ (right). Proteins were visualized by silver staining.

**Extended Data Fig. 8.**
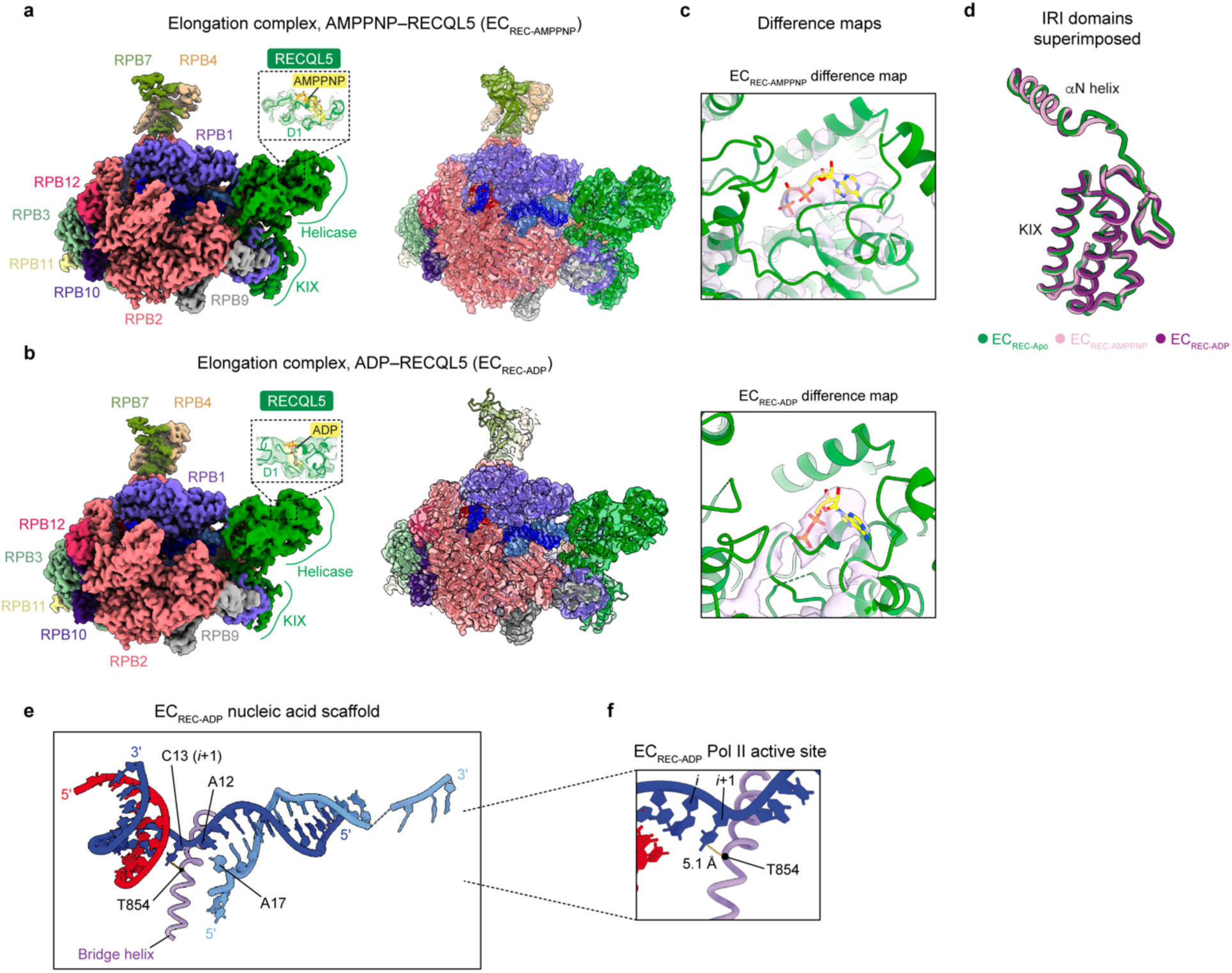
| Structures of ECs with nucleotide-bound RECQL5 helicase. **a**. Cryo-EM map (left) and cross-section with fitted atomic model (right) of Pol II EC bound to RECQL5_1– 620_ AMPPNP (EC_REC-AMPPNP_). Pol II subunits are colored as indicated by the labels, and nucleic acids are colored as in Fig. 1b. The inset shows the map and model around the helicase nucleotide binding site. **b.** Cryo-EM map (left) and cross-section with fitted atomic model (right) of Pol II EC bound to RECQL5_1– 620_ ADP (EC_REC-ADP_). Colors are the same as in **a**. The inset shows the map and model around the helicase nucleotide binding site. **c.** Difference maps for EC_REC-AMPPNP_ (top) and EC_REC-ADP_ (bottom). View shows the RECQL5 helicase nucleotide binding site. The helicase is depicted in green, and the modeled nucleotide in yellow. Details in **Methods**. **d.** Superposition of the RECQL5 IRI modules (αN helix and KIX domain) for the EC_REC-Apo_ (green), EC_REC-AMPPNP_ (pink), and EC_REC-ADP_ (purple) structures. **e.** Model of the nucleic acid scaffold and Pol II bridge helix in the EC_REC-ADP_ structure. Important nucleotides are highlighted. **f.** Close up of the EC_REC-ADP_ Pol II active site shown in **e** highlighting the distance between the *i*+1 site nucleotide base and RPB1 T854 in the bridge helix.

**Extended Data Fig. 9.**
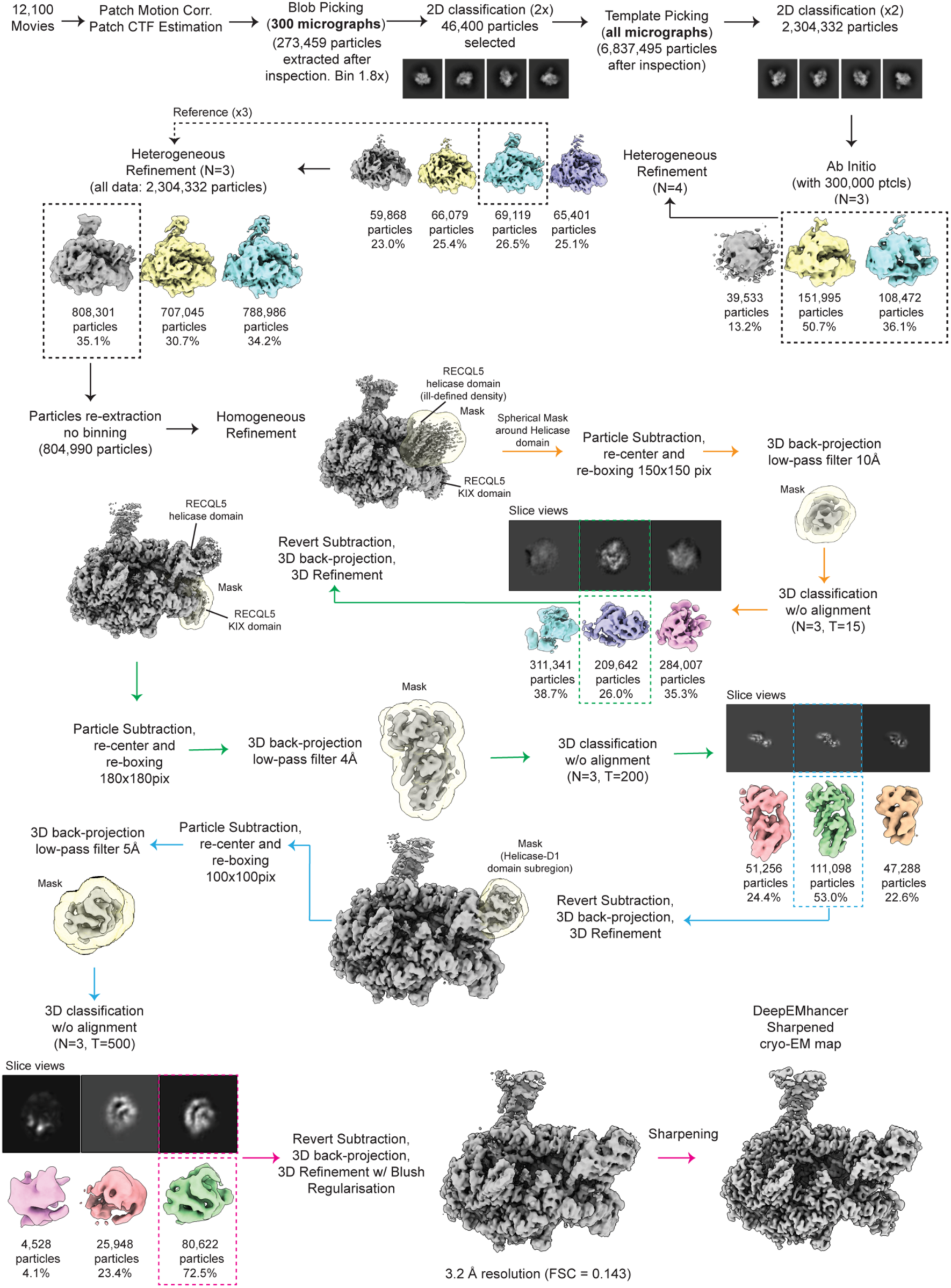
| Cryo-EM processing workflow for EC_REC-AMPPNP_.

**Extended Data Fig. 10.**
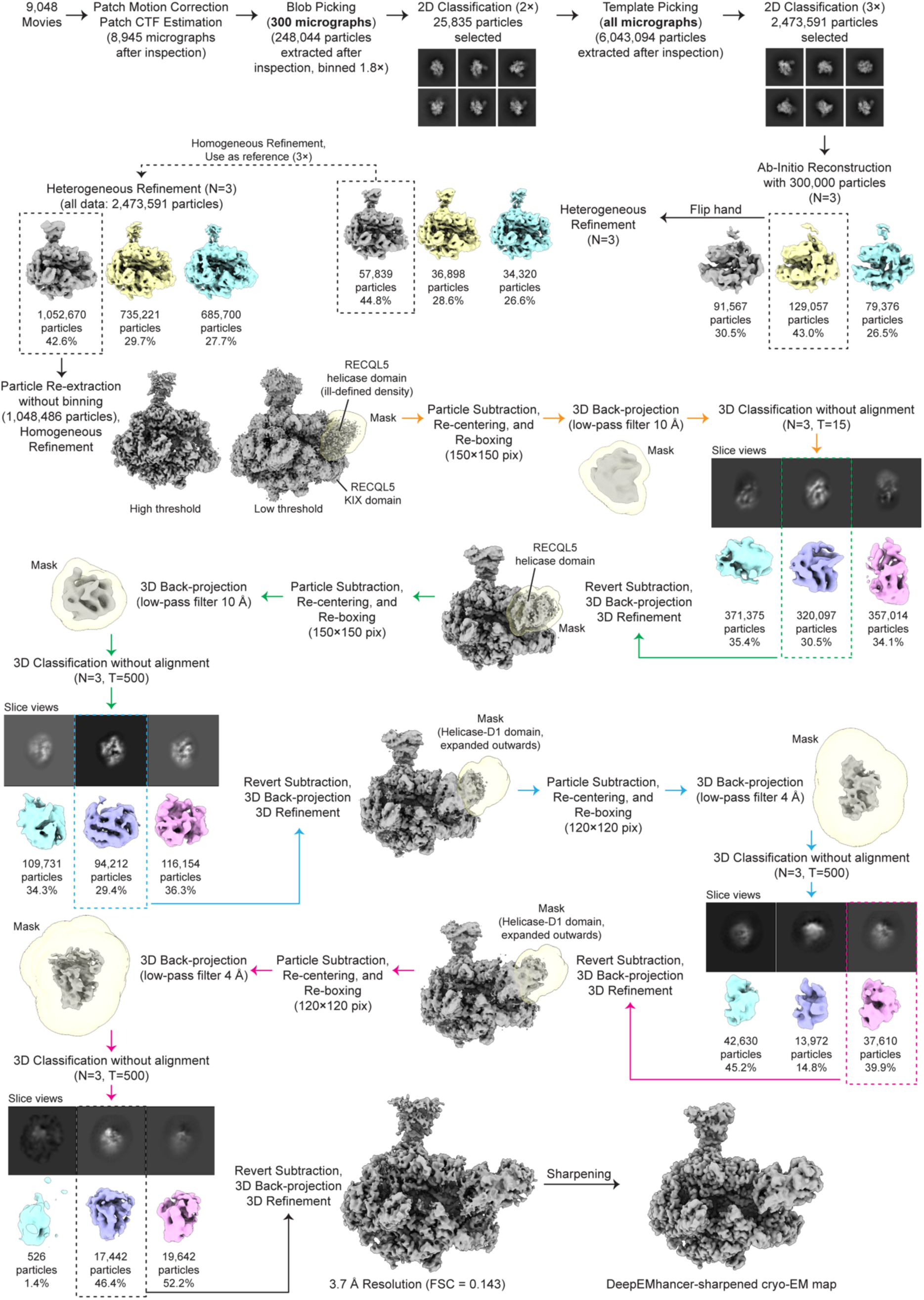
| Cryo-EM processing workflow for EC_REC-ADP_.

## Methods

### Protein purification

RECQL5 purification was as described in our previous study^22^. Briefly, GST-tagged RECQL5_1–620_ was expressed recombinantly in *E. coli* BL21-CodonPlus (DE3)-RIL cells (Stratagene) for 16 hours at 18° C, and the cells were lysed with a cell disrupter (Avestin). The clarified lysate was loaded onto a Glutathione-Sepharose 4 Fast Flow column equilibrated in RECQL5 buffer (20 mM Tris, pH 8.0, 150 mM NaCl, 10% glycerol, 1 mM dithiothreitol (DTT)), and eluted with a linear gradient to 20 mM glutathione. The GST tag was cut with PreScission protease during extensive dialysis for 14 h. The protein was subsequently purified using HiTrap Heparin HP and Superdex 200 columns (Cytiva) in RECQL5 buffer.

Human Pol II was purified according to a published protocol^34^. HeLa nuclei (114 l culture) was ground under liquid nitrogen using a mortar and pestle and then slowly resuspended in buffer A (50 mM Tris-HCl, pH 7.9 cold, 5 mM MgCl_2_, 0.5 mM ethylenediaminetetraacetic acid (EDTA), 25% glycerol, 5 mM DTT, 1 mM sodium metabisulfite, 1 mM phenylmethylsulfonyl fluoride (PMSF), supplemented with complete EDTA-free protease inhibitor cocktail (Roche)). After sonicating the resuspension for 2 min with stirring, (NH_4_)_2_SO_4_ was added to a final concentration of 0.3 M. The mixture was further sonicated to reduce viscosity and then clarified by centrifugation (40,000 rpm, 90 min, 4 °C, Ti45 rotor). The supernatant was adjusted to the conductivity of 0.1 M (NH_4_)_2_SO_4_ buffer through slow addition of buffer A. Then, a 42% (NH_4_)_2_SO_4_ cut was used to precipitate Pol II, followed by centrifugation (30,000 rpm, 30 min, 4 °C, Ti45 rotor). The precipitate was resuspended in buffer B (50 mM Tris-HCl, pH 7.9 cold, 0.1 mM EDTA, 25% glycerol, 2 mM DTT, 0.1 mM PMSF) with the concentration of (NH_4_)_2_SO_4_ adjusted to 0.15 M. The sample was applied to a DEAE52 column, which was washed with 3 column volumes of buffer B with 0.15 M (NH_4_)_2_SO_4_ before elution with buffer B with 0.4 M (NH_4_)_2_SO_4_. Protein-containing fractions were pooled and adjusted by dialysis to the conductivity of 0.2 M (NH_4_)_2_SO_4_ buffer, supplemented with 0.1% NP-40 substitute, and immunoprecipitated overnight at 4 °C using anti-RPB1 antibody (clone 8WG16, Biolegend) crosslinked to Protein G Sepharose Fast Flow resin (Cytiva). The resin was washed three times with buffer C (25 mM HEPES, pH 7.9 cold, 0.2 mM EDTA, 10% glycerol, 2 mM DTT, 0.1 mM PMSF, 0.05% NP-40 substitute) with 0.5 M (NH_4_)_2_SO_4_, followed by two washes with buffer C with 0.2 M (NH_4_)_2_SO_4_. Pol II was eluted through four sequential incubations with buffer C supplemented with 0.23 M (NH_4_)_2_SO_4_ and 1 mg ml^−1^ RPB1 tri-heptapeptide repeat peptide (sequence: (YSPTSPS)_3_). Concentrated eluate was flash-frozen in liquid N_2_. An SDS-PAGE gel of purified proteins is shown in **Extended Data Fig. 7b**.

### Preparation of EC_REC-Apo_ complex

The purification and crosslinking of the apo complex (Pol II bound to the nucleic acid scaffold and RECQL5_1–620_ D157A with no nucleotide) was described in our previous paper^22^. Briefly, Pol II was incubated first with a ten-fold molar excess of the nucleic acid scaffold and then with a ten-fold molar excess of RECQL5 while immobilized to the anti-RPB1–Protein G resin during the Pol II purification procedure (see above). The resin was washed and the EC_REC-Apo_ complex was eluted with RPB1 tri-heptapeptide repeat peptide as usual. Then, the complex was diluted with transcription buffer (20 mM HEPES, pH 8.0, 4 mM MgCl_2_, 50 mM KCl, 0.05% NP-40 substitute, 1 mM Tris-(2-carboxyethyl)phosphine (TCEP)) and crosslinked with 0.02% glutaraldehyde for 10 min, followed by quenching with 100 mM Tris. Complex was then aliquoted and flash-frozen with liquid N_2_.

### Preparation of EC_REC-AMPPNP_ and EC_REC-ADP_ complexes

Nucleotide-bound complexes were assembled and purified using a pulldown strategy (**Extended Data Fig. 7a**). First, tDNA (5’-CTCAAGTACTTACGCCTGGTCATTACTA-3’) and RNA (5’-UAUAUGCAUAAAGACCAGGC-3’) were annealed by incubating at 90 °C for 5 min and then cooling to 4 °C at a rate of 0.2 °C s^−1^. Pol II (diluted from 513 nM stock) was mixed with the tDNA– RNA hybrid and incubated at room temperature for 20 min. Then, desthiobiotinylated ntDNA (5’-/5deSBioTEG/TAGTAAACTAGTATTGAAAGTACTTGAGCTTAGACAGCATGTC-3’) was added, and the mixture was incubated at room temperature for 20 min. All oligonucleotides were purchased from Integrated DNA Technologies. Finally, RECQL5_1–620_ and nucleotide were added, and the mixture was incubated at room temperature for 20 min. Because D157 stabilizes bound ADP through water-mediated interactions with the coordinated Mg^2+^ ion^32^, we employed wild-type RECQL5 for these studies instead of the D157A mutant. The final mixture contained 350 nM Pol II, 350 nM nucleic acid scaffold, 7 μM RECQL5, and 1 mM AMPPNP or ADP, diluted in transcription buffer. After the incubation, Dynabeads MyOne Streptavidin T1 was added (6.25 μl beads per 12.5 μl input) and the mixture was incubated at room temperature for 75 min. The beads were washed twice with transcription buffer containing 1 mM of AMPPNP or ADP, as appropriate. Then, the complex was eluted by incubating the beads twice with elution buffer (20 mM HEPES, pH 8.0, 4 mM MgCl_2_, 50 mM KCl, 0.05% NP-40 substitute, 1 mM TCEP, 5 mM biotin, 3% trehalose, 1 mM AMPPNP or ADP) at 37 °C for 15 min. The eluted complex was crosslinked with 0.02% glutaraldehyde at room temperature for 10 min and quenched by incubating with 100 mM Tris-HCl, pH 8.0, at room temperature for 5 min. The crosslinked complexes were used to prepare cryo-EM grids on the same day.

### Cryo-EM sample preparation

Cryo-EM specimens were deposited on graphene oxide (GO)-coated Quantifoil grids (1.2/1.3 300-mesh, carbon-on-gold). To prepare the GO-coated grids, we adapted our previously reported protocol^35^. Grids were cleaned with two drops of chloroform each, glow-discharged using a Tergeo-EM plasma cleaner (PIE Scientific), incubated for 2 min with 1 mg ml^−1^ polyethylenimine (Polysciences) in 25 mM HEPES at pH 7.9, washed twice with H_2_O, and air-dried. Then, grids were incubated for 2 min with 0.2 mg ml^−1^ GO stock solution, washed twice with H_2_O, and air-dried. To prepare the GO stock solution, we diluted GO in 1:2 methanol:H_2_O (v:v), sonicated the mixture, centrifuged at 4,000*g* for 10 min (to remove small GO sheets), resuspended the pellet in 1:2 methanol:H_2_O (v:v), further sonicated the mixture, and finally collected the supernatant after centrifugation at 1,000*g* for 1 min (to remove GO aggregates). We found that a 1:2 methanol:H_2_O (v:v) solution facilitated the deposition of a continuous GO layer on the grid. Grids were either used on the same day or saved and gently glow-discharged before use. Onto each grid, 3.5 μl of sample was deposited, followed by incubation for 30 s at 22°C with 100% humidity in a Vitrobot Mark IV (Thermo Fisher Scientific). Then, the grid was blotted for 10 s with a blot force of 10 and vitrified by plunging into liquid ethane with a liquid N_2_ bath.

### Cryo-EM data collection

All cryo-EM data were acquired as dose-fractionated movies with a 300 kV Titan Krios G3 cryo-electron microscope (Thermo Fisher Scientific) using a K3 direct electron detector (Gatan). A total exposure dose of 50 *e*^−^/Å^2^ fractionated across 50 frames was used during movie frame recording, with defocus values ranging from approximately −0.8 µm to −1.8 µm. All data collection processes were automatically controlled using SerialEM^36^ and parameters are summarized in **Table 1**.

For the EC_REC-Apo_ sample, 6,190 movies (Dataset 1) were collected in super-resolution counting mode at 81,000× magnification using correlated double sampling (CDS) and a super-resolution pixel size of 0.525 Å pixel^−1^. From Dataset 1, the EC_REC-Apo_, EC_Free_ and EC_REC-Apo (IRI-focused)_ structures were produced (described below).

With the aim of increasing the number of movies acquired per hour, for the EC_REC-AMPPNP_ and EC_REC-ADP_ complexes, data were acquired in the same instrument as described above but using a non-CDS/non-super-resolution mode, at 81,000× magnification with a physical pixel size of 1.048 Å pixel^−1^. For EC_REC-AMPPNP_ and EC_REC-ADP_, a total of 12,100 (Dataset 2) and 9,048 movies (Dataset 3) were collected, respectively.

### Cryo-EM image processing

Data processing of all the cryo-EM datasets was performed using cryoSPARC v4.5.3^37,38^ and RELION5^39,40^ software as detailed in **Extended Data Figs. 1, 3, 5, 9, and 10**. For simplicity, we will describe in detail the data analysis workflow followed for Dataset 1, which resulted in the elucidation of the EC_Free_, EC_REC-Apo_, and EC_REC-Apo (IRI-focused)_ structures (**Extended Data Figs. 1, 3, and 5**, respectively). We note that the analyses for Dataset 2 and Dataset 3 (EC_REC-AMPPNP_ and EC_REC-ADP_ structures, respectively) were performed following similar workflows as for EC_REC-Apo_.

#### Initial Dataset 1 processing

For Dataset 1, the 6,190 movie frames collected were aligned using Patch Motion Correction within cryoSPARC^37,38^. Then, defocus estimation and contrast transfer function (CTF) fitting were performed using Patch CTF Estimation. In the corrected micrographs, we could readily observe particles with the size and features expected for Pol II ECs (**Extended Data Fig. 2a**).

A preliminary round of data processing was performed on 100 micrographs randomly selected, where particles were picked using the Blob Picker algorithm. Later, multiple rounds of 2D classification/particle selection cycles were carried out to obtain suitable 2D templates for the following Template Picker job on all micrographs. Three subsequent rounds of 2D classification/particle selection cycles resulted in a set of 2,240,800 particles, from which 300,000 particles were randomly selected to perform an *ab initio* 3D reconstruction (n = 3). Two out of three 3D *ab initio* maps were used as references to run a Heterogeneous Refinement using the full particle set (n = 4). From these classes, one particular class, containing 39.1% of the population (875,262 particles), showed defined structural features, while the other classes displayed broken complexes and/or poor low-resolution reconstructions. The particles corresponding to the best class were re-extracted using a box size of 320 pixels × 320 pixels (without binning), resulting in 871,524 particles (duplicate particles removed), and further subjected to a Homogeneous Refinement job, obtaining a 3D reconstruction at 2.4 Å overall resolution (FSC = 0.143). At low-threshold levels, two fuzzy regions appeared next to the EC, resembling quite well the positioning of the RECQL5 helicase (region A) and KIX domains (region B) observed in the low-resolution cryo-EM structure of this complex reported previously by our group^22^. These ill-defined densities suggested a large degree of local heterogeneity or partial occupancy of RECQL5, which was quite difficult to sort out by standard classification methods. Therefore, we implemented the data analysis pipeline detailed below.

#### EC_REC-Apo_ processing

The 871,524 particles from the Homogeneous Refinement job in cryoSPARC were exported to RELION^39,40^ and subjected to 3D Refinement. The RECQL5 helicase domain appeared significantly less stable than the KIX domain area. Therefore, we aimed first to resolve the local heterogeneity in the helicase domain (region A).

Using the Volume Segmentation tool in ChimeraX^41,42^ and the Mask Creation job in RELION, we generated a binary mask involving the region assigned to the RECQL5 helicase domain. We then performed a Particle Subtraction job to keep the signal inside the mask, at the same time re-centering the subtracted particles on the mask and re-boxing them to a box size of 180 pixels × 180 pixels. Then, using the relion_reconstruct program, we back-projected the subtracted particles to generate a lowpass filtered 3D reconstruction to be used as a 3D reference for the next 3D classification job. This 3D classification job was performed without alignment, applying a contoured mask, generating four (n = 4) classes, and using a T value of 15 and the Blush Regularization option activated. One of the four classes, containing 23.7% of the population (206,338 particles) and displaying better defined features, was selected and subjected to subtraction reversion in order to recover the full particle information. The reverted particles were then back-projected using the relion_reconstruct program to generate a new 3D reference and then subjected to 3D Refinement resulting in a reconstruction at 3.3 Å overall resolution. In this map, the RECQL5 helicase domain showed significant improvement (interestingly, the KIX domain density improved as well), and defined structural features started to become apparent. We followed up with an additional round of this strategy, but in the second round of 3D classification without alignment the T value was increased to 500. We suspected that a larger T value would be helpful since more relative weight will be considered on the actual experimental data (particles) over the prior, along the classification cycles. One major class, accounting for 42.1% of the population (86,831 particles), and displaying clear secondary structure features, was selected and subjected to subtraction reversion, back-projection and 3D Refinement (same as in the first cycle). The resulting reconstruction displayed, overall, a well-defined RECQL5, although some fuzziness was still observed for the helicase D1 subdomain (for orientation, see **Fig. 1a**). Therefore, we performed two additional rounds of this particle subtraction/3D classification/subtraction reversion/3D refinement cycle dedicated to improving the helicase D1 subdomain region. To this aim, different combination of particle re-boxing sizes and T values were used, since these parameters needed to be adapted to the smaller region under analysis. Ultimately, 24,323 particles were used to obtain the final cryo-EM reconstruction of EC_REC-Apo_ at 3.2 Å overall resolution (FSC=0.143, **Extended Data Fig. 2c**). In this map, the Pol II EC core has the highest local resolution (Fig Supp X3c), while the fully visible RECQL5 helicase and KIX domain regions have local resolutions ranging between 3.7 Å and 5.9 Å (**Extended Data Fig. 2c**). The final cryo-EM reconstruction was post-processed using the DeepEMhancer sharpening program^43^.

The data analysis approach described above was used for EC_REC-Apo_, as well as the EC_REC-AMPPNP_, and EC_REC-ADP_ and enabled us to improve the helicase domain resolution within the context of the full complex. For the purpose of elucidating molecular interactions between RECQL5 and the Pol II EC, this workflow worked better than standard approaches like focused classification/focused refinement only, which only result in an improved helicase domain map isolated from the rest of the EC. It allowed us to map the full RECQL5_1–620_ construct and describe the extent of its interactions with different regions the Pol II EC (**Fig. 2**).

#### EC_REC-Apo (IRI-focused)_ processing

We employed a similar workflow to further improve the local resolution of the RECQL5 KIX domain region (region B). Starting over from the 871,524 particles exported to RELION, we subjected the particles to 3D Refinement and then adapted our previous data analysis approach to solve the local heterogeneity in this area (**Extended Data Fig. 5**).

By rigid-body fitting both initial Pol II EC coordinates^27^ (PDB 5FLM) and a RECQL5 model predicted by AlphaFold 3^44^ into the EC_REC-Apo_ cryo-EM map, we observed that the RECQL5 IRI module (harboring the αN helix and KIX domain) is positioned to interact with the lower jaw of the Pol II RPB1 subunit. Therefore, using the Volume Segmentation tool in ChimeraX^41,42^ and the Mask Creation job in RELION^39,40^, we generated a binary mask involving both the RPB1 lower jaw and the RECQL5 IRI module regions. We then performed a Particle Subtraction job to keep the signal inside the mask, re-centering of the subtracted particles on the mask and re-boxing them to a box size of 180 pixels × 180 pixels. Then, using the relion_reconstruct program, we back-projected the subtracted particles to generate its own low-pass filtered 3D reconstruction to be used as a 3D reference for the next 3D classification job. This 3D classification job was performed without alignment, applying a contoured mask, generating four (n = 4) classes, and using a T value equal to 20 together with the Blush Regularization option activated. One of these four classes, harboring 29.0% of the population (253,055 particles), displayed better features clearly observed both in the map and in the slice view representation as well. This class population was selected and subjected to subtraction reversion in order to recover the full particle information. The reverted particles were then back-projected using the relion_reconstruct program and subjected to 3D Refinement obtaining a reconstruction at 3.2 Å overall resolution. In this 3D map, a clear improvement of the RECQL5 KIX domain region is observed. Thus, we decided to perform one additional round of this particle subtraction/3D classification/subtraction reversion/3D Refinement cycle. Ultimately, 103,215 particles were used to obtain the final cryo-EM reconstruction of the EC_REC-Apo (IRI-focused)_ at 2.8 Å overall resolution (FSC=0.143, **Extended Data Fig. 2d**). In this map, the region of interest involving the RECQL5 KIX domain is fully visible and displayed a local resolution range of about 3.2–4.0 Å (**Extended Data Fig. 2d**), a major improvement compared to the EC_REC-Apo_ structure. This final cryo-EM map was then post-processed using the DeepEMhancer sharpening program^43^.

#### EC_Free_ processing

We also wanted to investigate the structure of the free stalled Pol II EC in the absence of RECQL5 (EC_Free_). As mentioned above in the EC_REC-Apo_ data analysis section, the 2.4 Å resolution cryo-EM map obtained from Homogeneous Refinement of 871,524 un-binned particles (e.g., prior to RELION processing) showed two fuzzy regions next to the EC corresponding to the RECQL5 helicase (region A) and KIX domains (region B) (**Extended Data Fig. 3**). These ill-defined densities suggested a large degree of local heterogeneity or partial occupancy of RECQL5. Therefore, in order to take advantage of this possible RECQL5 partial occupancy, we performed global 3D classification without alignment in cryoSPARC (**Extended Data Fig. 1**). From the four 3D classes generated, one class harboring 24.3% of the total population (212,196 particles) showed no density attributable to any RECQL5 domain and was therefore recognized as a free EC. This class population was selected and subjected to Homogeneous Refinement, resulting in a 2.6 Å resolution (FSC = 0.143) cryo-EM map. Then, an additional round of 2D classification was used to remove low-resolution particles and remaining contaminants, resulting in a set of 174,428 particles. Lastly, Non-uniform Refinement was used to obtain the final EC_Free_ cryo-EM structure at 2.4 Å overall resolution (FSC = 0.143, **Extended Data Fig. 2b**). This final cryo-EM map was then post-processed using the DeepEMhancer sharpening program^43^.

#### Dataset 2 and Dataset 3 processing

The image processing corresponding to Dataset 2 and Dataset 3 was performed following the same approach used for Dataset 1 to obtain the EC_REC-Apo_ structure. As detailed in **Extended Data Figs. 9 and 10**, a total of 80,622 and 17,442 particles were used to obtain the final structures of EC_REC-AMPPNP_ (3.2 Å overall resolution, FSC = 0.143) and EC_REC-ADP_ (3.7 Å overall resolution, FSC = 0.143), respectively. Both maps were then post-processed independently using the DeepEMhancer sharpening program^43^.

### Model Building, refinement, and validation

For the EC_Free_, EC_REC-Apo_, EC_REC-AMPPNP_, and EC_REC-ADP_ structures, the initial coordinates of the EC were obtained by rigid-body fitting the atomic model of the transcribing mammalian Pol II^27^ (PDB 5FLM) into the corresponding post-processed maps using ChimeraX^41,42^. For RECQL5, the initial coordinates were obtained from different sources. For the helicase and RQC domains, the initial model was taken from the X-ray structure of the human RECQL5 in apo form^32^ (PDB 5LB8), while for the αN helix and KIX domain, the atomic model was predicted using AlphaFold 3^44^. These RECQL5 model regions were then semi-automatically docked and rigid-body fitted into the corresponding sharpened map.

For EC_REC-Apo (IRI-focused)_, the transcribing mammalian Pol II^27^ (PDB 5FLM) model and the predicted coordinates for the RECQL5 IRI module (αN helix and KIX domain) were both rigid-body fitted into the sharpened map. For subsequent model building and refinement, we only kept the coordinates corresponding to the Pol II RPB1 lower jaw (residues 1162–1305) and the RECQL5 IRI module (harboring the αN helix and KIX domain, residues 498–620), since these were the interacting regions of interest for this structure.

For each complex, the models were then iteratively rebuilt in *Coot*^45^ and refined using the real space refinement program in *Phenix*^46^. All validation and refinement statistics are shown in **Table 1**.

All plots of FSC_map vs map_ and FSC_map vs model_ were obtained using a python script developed in-house. The FSCs_map vs map_ were obtained from the half-maps for each structure considering an applied contoured mask. The FSCs_map vs model_ were obtained by running a validation job in *Phenix* of the corresponding final refined atomic model against the unsharpened full map.

### Structural visualization and interpretation

All the structural comparisons and superpositions, as well as rotation, root-mean-square-deviation (RMSD), and distance measurements were performed in ChimeraX^41,42^. The difference maps showed in **Extended Data Fig. 8c** were generated in ChimeraX as follows: first, we created a density map from the coordinates corresponding to Pol II EC and RECQL5 only, without considering the nucleotide bound; then, this map was subtracted from the corresponding full cryo-EM map for EC_REC-AMPPNP_ or EC_REC-ADP_. Overall, all main figures show the sharpened cryo-EM maps and the final refined atomic models unless otherwise specified.

## Data Availability

The cryo-EM density maps and their respective atomic coordinate files were deposited to the Electron Microscopy Data Bank (EMDB) and Protein Data Bank (PDB) under the following accession codes: EMD-48071 and PDB 9EHZ (EC_Free_), EMD-48073 and PDB 9EI1 (EC_REC-Apo_), EMD-48074 and PDB 9EI2 (EC_REC-Apo (IRI Focused)_), EMD-48075 and PDB 9EI3 (EC_REC-AMPPNP_) and, EMD-48076 and PDB 9EI4 (EC_REC-ADP_). Other data are available from the corresponding author on request.

